# Patient-Derived Surgical samples reveal patterns of glioblastoma infiltration and tumor microenvironment at the tumor margin

**DOI:** 10.1101/2025.03.09.633849

**Authors:** Olaya de Dios, Juan Romero, María de los Ángeles Ramírez-González, Beatriz Herranz, Alicia Avis, Ana Ramos, Juan Manuel Sepúlveda-Sánchez, Ricardo Gargini, Berta Segura-Collar, Gabriel Velilla, Bárbara Meléndez, Pedro González, Luis Jiménez-Roldán, Guillermo García-Posadas, Aurelio Hernández-Laín, Ángel Pérez-Núñez, Pilar Sánchez-Gómez

## Abstract

**Background:** Glioblastoma (GBM) is a highly aggressive cancer with near-universal recurrence, often due to residual tumor cells that persist after aggressive standard of care treatment. This study aimed to characterize tumor infiltration and microenvironment in the GBM periphery.

**Methods:** We prospectively collected 161 radiologically guided biopsies from 45 GBM patients and conducted an immunohistochemical analysis. We also integrated single-cell RNA sequencing data to select specific markers for immune, glial, and vascular cells. We measured the expression of these genes in samples from contrast-enhancing (CE) and non-enhancing (nCE) tumor areas, vasogenic edema, and radiologically normal tissue. Correlations with resection extent and clinical outcomes were evaluated.

**Results:** nCE biopsies exhibited neoplastic features similar to those of the tumor core. However, tumor infiltration was also found in regions classified radiologically as edema, particularly in elderly patients. We found important differences in the composition of the peripheral microenvironment between male and female GBM patients. Prognostic associations with specific cell types, such as myeloid cells, showed intertumor heterogeneity, with variations depending on patient sex, age and extent of resection. Furthermore, in our cohort, minimal residual CE tumor following surgery was associated with significantly poorer patient survival.

**Conclusions:** The GBM periphery includes regions of active tumor growth that are visible on MRI, as well as infiltrated areas that resemble edema radiologically. Tumor infiltration and microenvironmental features are influenced by patient sex and age, which has major implications for recurrence rates, highlighting the need to tailor surgical and therapeutic strategies based on tumor biology and patient subgroup.

**Key points:** nCE areas show similar neoplastic traits to the CE tumor.

GBM infiltrates edema tissue, predominantly in older adult patients.

Prognostic value of the peritumoral phenotype depends on resection extent and patient age/sex.

**Importance of the study:** This study shows that the impact of peripheral and core cellular features on prognosis differs between patients who undergo complete versus incomplete CE tumor resection. Our results suggest a paradigm shift in the classification and management of these patients, encouraging the inclusion of detailed post-surgical MRI analyses to guide the design of future clinical trials according to the nature and extent of the residual disease. Furthermore, our data confirm the presence of tumorigenic features in non-enhancing areas, supporting the benefits of supratotal resections according to the new RANO classification. Our findings also underscore the need to refine surgical and therapeutic strategies based on a more detailed understanding of the tumor microenvironment beyond the GBM core. This understanding may help identify novel targets for more effective and personalized GBM therapies.

## Introduction

Glioblastoma (GBM) is the most common and lethal central nervous system (CNS) tumor, with a five-year survival rate of only 5% [1]. Magnetic resonance imaging (MRI) is the gold standard for diagnosing and characterizing these tumors, which appear as intra-axial infiltrative lesions with ill-defined margins that blend into the surrounding brain tissue. GBM typically presents as a contrast-enhancing (CE) lesion that is sometimes heterogeneous due to intra-tumoral necrosis, cysts, or hemorrhage. The core is surrounded by a hyperintense area in T2-FLAIR (fluid-attenuated inversion recovery) sequences.

Although gliomas rarely metastasize, they infiltrate the brain parenchyma. While invasiveness is not exclusive to the deadliest GBM, it is a key driver of treatment resistance in these tumors. This feature explains why GBM invariably recurs, typically at the edge of the resection cavity, even after complete resection of the CE tumor followed by radiotherapy targeting the tumor bed and systemic treatment with temozolomide [2]. Recent studies suggest that infiltrative GBM regions can be detected using FLAIR sequences, where the non-enhancing (nCE) tumor appears as lesions with mild-to-high signal intensity lesions [3, 4]. However, distinguishing nCE tumor from edema and gliosis, which also present as high FLAIR signals, remains challenging and depends largely on subjective radiological criteria. Furthermore, some GBM cases lack detectable nCE tumor, indicating that infiltrative tumor areas are not always visible on conventional MRI.

The concept of supratotal resection has gained traction in recent years. This approach aims to remove as much of the tumor as possible, including both CE and nCE, as long as it can be done safely without causing significant neurological deficits. Several studies have reported a significant increase in overall survival (OS) for GBM patients, averaging an additional six months, an outcome unmatched by any other oncological treatment to date [5, 6]. In light of these findings, new Response Assessment in Neuro-Oncology (RANO) criteria have been proposed to classify the extent of surgical resection [7], and clinical trials are underway to validate the survival benefits of these extended resections [8].

These results underscore the importance of characterizing the hyperintense FLAIR region in the periphery of GBM tumors. This area has been shown to contain a mixture of tumor cells, immune system cells, reactive astrocytes, and cells from the oligodendrocyte lineage, within a vascular context that differs significantly from that of the tumor core [9-12]. However, despite these advancements, large-scale studies investigating variations between distinct radiological regions or differences among patient subgroups remain scarce. Furthermore, none of these studies have correlated their findings with GBM progression and outcomes.

In this study, we prospectively generated a cohort of 161 radiologically guided biopsies from the enhancing nodule region and its periphery. We also assessed the extent of resection in the 45 patients included in the study. To evaluate neoplastic infiltration in the samples, we performed an immunohistochemical (IHC) analysis of proliferative cells. Additionally, we used publicly available single-cell RNA sequencing (scRNAseq) data to identify genes specific to different cell types in the brain microenvironment. Our IHC study confirmed that nCE tumor areas contain a significant number of proliferating cells, with a density similar to that of the tumor core. However, we also identified tumor features in regions radiologically classified as vasogenic edema and in areas with no radiological alterations, particularly in older adult patients. Furthermore, unsupervised clustering of differential gene expression analysis distinguished the most pathogenic samples based on the enrichment of proliferative markers and the absence of normal endothelial and oligodendrocyte expression. Distinct cellular markers in the tumor core and peripheral edema, particularly those associated with immune cells, astrocytes and proliferation, were linked to GBM prognosis. These features exhibited different trends in patients who underwent complete versus incomplete resection of the CE tumor. Additionally, significant sex and age-based differences in the cellular characteristics of the GBM periphery were identified that were associated with distinct prognostic outcomes. Our results suggest that the nature of the infiltrative zone depends on the tumor core phenotype, as well as on age and sex. Moreover, therapeutic strategies targeting residual GBM disease must carefully consider the extent of resection and patient subgroup characteristics.

## Methods

The workflow followed in this manuscript is illustrated in Supplementary data S1A.

### Human samples

A prospective study was conducted from April 2021 through April 2024 on all adult patients undergoing surgery for a novel diagnosis of high-grade glioma, excluding those receiving stereotactic biopsies or cases in which obtaining multiple biopsies was deemed to pose unnecessary risk. We only included patients with a confirmed diagnosis of GBM with wild-type *isocitrate dehydrogenase* (IDH) 1 and 2 genes. Preoperative MRI scans were used to determine tumor volume and location and to select the tumor regions for biopsy collection, prioritizing the most extensive safe tumor resection for each patient. Screenshots were taken during surgery to confirm the sample locations.

At least two radiologists reviewed the surgical images and categorized the biopsies into CE tumor, nCE tumor, vasogenic edema, or radiologically normal regions. Biopsies obtained in areas of FLAIR hyperintensity without contrast enhancement were subcategorized into: non-enhancing tumour and vasogenic edema. We used observational criteria to differentiate the samples in these two subcategories. Our neuroradiology team considered as nCE: all those biopsies obtained in areas with nodular or expansive behavior, or with cortical thickening, and/or with lower hyperintensity in FLAIR sequence than vasogenic edema. The Apparent Diffusion Coefficient (ADC) value of these samples was measured with Olea Sphere © and was less than 1 x 10^-3^ mm2/s in all our cases. To label the samples as vasogenic edema we considered all those areas with a very high signal on the FLAIR sequences with digitiform pattern that were not included in the nCE tumor category. Most of these samples had ADC values between 1 x 10^-3^ and 2 x 10^-3^ mm2/s. Postoperative MRI scans (obtained within ≤72 hours after surgery whenever possible) were analyzed by radiologists to assess the extent of resection for each patient. Histopathological examination and sequencing allowed us to exclude all IDH mutant (IDHmut) cases, as well as CE tumor biopsies with high necrotic content. A detailed list of patients and collected samples is provided in Supplementary data S1B.

### Molecular characterization of the tumors

Biopsies taken during surgery (and before treatment) were stored in RNAlater (frozen) or fixed in PFA and embedded in paraffin (FFPE). DNA was extracted from FFPE tumor tissues using the Maxwell RSC DNA FFPE Kit (Promega) and quantified using Qubit 2.0 Fluorometer (Thermo Fisher Sci.). We used a custom Ampliseq (PCR-based) gene-targeted next-generation sequencing (NGS) panel to analyze common mutations: TERTp, 1p/19q codeletion, EGFR/EGFRviii/PDGFRA amplifications, and CDKN2A/PTEN deletions [13]. Ampliseq libraries were manually prepared using Ion AmpliSeq Library Kit Plus and sequenced on the Ion Torrent Genexus Integrated sequencer (Thermo Fisher). MGMT methylation status was assessed with the Pyromark Q24 kit (QIAGEN).

### RNA Extraction and qRT-PCR

RNA was extracted using TRIzol (Invitrogen), following the protocol detailed in Supplementary data S1D. 1 μg of RNA was reverse transcribed using the PrimeScript RT Reagent Kit (Takara). qRT-PCR was performed using SYBR Premix (Takara) and gene-specific primers on the LightCycler 480 (Roche Diagnostics) (Supplementary data S1C). *GAPDH, HPRT*, and *RPII* were tested as candidate housekeeping genes. After assessing expression consistency across these genes, *GAPDH* was selected for normalization. Reactions were run in technical duplicates, and no-template controls were included. Relative gene expression was calculated using the ΔΔCt method.

### Immunohistochemical (IHC) staining and quantification

MIB-1 IHC staining (Supplementary data S1C) was performed on 4 μm FPPE sections. Sslides were stained with hematoxylin and eosin to confirm the histological features. For chromogenic detection, slides were preheated for 30 minutes at 65°C, and the BOND RXm automated advanced staining system (Leica Biosystems) was programmed according to the specific staining protocol (Supplementary data S1E). Slides were scanned using the Hamamatsu NanoZoomer-SQ microscope, and quantification was performed with QuPath software using positive-cell detection method, which calculates the percentage of positive cells relative to negative cells within a consistently defined area for each stain (Supplementary data S1F).

### scRNA-seq datasets analysis

We analyzed our previously published integrated human single-cell atlas, which was generated by integrating publicly scRNA-seq datasets from normal brain, IDHmut gliomas (lower-grade gliomas, LGG), and GBM, as described previously [14] (Supplementary Figure S1) (Supplementary data S1C). We included non-tumoral, LGG and GBM samples so that the gene list could be potentially used will all glioma samples, although the results presented here are restricted to GBM. To validate the key genes in the tumor periphery, processed RDS files [15] were downloaded from public repositories. Data processing and analysis were conducted using the Seurat v4.3.0. Differentially expressed genes (DEGs) were determined with the Seurat functions *FindAllMarkers* (for Cell-Type Specific Markers) and *FindMarkers* (for condition-specific comparisons), requiring gene expression in at least 25% of cells and a minimum log fold-change of 0.25.

### Key Genes identification

Two different types of analysis were performed (Supplementary Figure S2).

#### Analysis 1

Identification of Key Genes from DEGs:

(1) Cell-Type Specific Markers from the Integrated Single-Cell Atlas. DEGs that are markers of each cell type, using the following parameter across clusters: ptc.2 ≤ 0.1 for most of the clusters, ptc.2<0.2 for Astrocytes-C10, and ptc.2 < 0.3 for endothelium, mural cells, and neurons (Supplementary data S2.A1). DEGs in Control brains vs. LGG Patients within each specific cluster (parameter settings: avg_log2FC =<-1) (Supplementary data S2.A2). (3) DEGs in Control brains vs. Newly Diagnosed GBM Patients (parameter settings: avg_log2FC =<-1) (Supplementary data S2.A3). (4) DEGs in Control brains vs. Recurrent GBM Patients. DEGs in homeostasis versus newly recurrent GBM patients (Supplementary data S2.A4) within each specific cluster (parameter settings: avg_log2FC =<-1).

Genes overlapping across these lists within each specific cell type (Supplementary data S2.A5) were defined as key gene candidates (Supplementary data S1G). They were further validated by comparing GTEx Brain-Cortex/Frontal Cortex versus TCGA LGG and GBM datasets (Supplementary data S1G – List 1). To assess significant expression differences, we applied Welch’s t-test (TCGA vs. GTEx) [16], identifying the most differentially expressed genes between normal brain tissue and glioma samples (Supplementary data S1G – List 2).

#### Analysis 2

Identification of key genes from conservative markers (Supplementary data S2.B2). We performed a Venn Diagram-based comparison between the significative conservative markers in the control brain and glioma samples, alongside cell-type specific markers identified in the Integrated Single-Cell Atlas (parameter settings: ptc.2 ≤ 0.1) (Supplementary data S2.B1) (Supplementary data S2.B3). It was applied when Analysis 1 did not yield strong markers for certain cell types, helping us to extract key markers for endothelial cells and oligodendrocytes.

Candidate genes from both analyses were finally candidate visualized in the integrated single-cell atlas and the Core/Peripheral database to select the most cell-type-specific genes (Supplementary data S3).

### Survival analysis

To assess the prognostic value of the expression of the markers in GBM we used the *The Human Protein Atlas* web tool [17] (proteinatlas.org). RNA sequencing (RNAseq) data came from the TCGA. The best cutoff algorithm was used to divide GBM cancer patients into high- or low-expression groups (Supplementary data S1G).

### Cell Type RNA expression

To confirm cell-type-specific genes of the markers we employ the *The Human Protein Atlas* web tool [17]. The RNA specificity category is based on mRNA expression levels in the analyzed cell types based on scRNA-seq data from normal tissues (Supplementary data S1G).

### Statistics

All analyses were conducted using R (v.4x.) and GraphPad Prism (10.1.2). Prior to analysis, continuous variables were log10-transformed and standardized (z-score). Normality was assessed using the Shapiro-Wilk test, and due to the non-normal distributions and the small subgroup sizes, non-parametric tests were applied throughout. For comparisons between two groups, the Wilcoxon rank-sum test (Mann-Whitney U test) was used. For comparisons across multiple groups, the Kruskal-Wallis test followed by Dunn’s post hoc test was performed. Correlations were evaluated using Spearman’s coefficients (ρ). Survival analysis was conducted with Kaplan-Meier curves and log-rank test.

Unsupervised clustering of tumor samples was performed (ConsensusClusterPlus, v.1.64.0) based on the expression matrix of 45 genes. Clustering stability was evaluated across a range of cluster numbers (K = 2 to 10) using hierarchical clustering with Ward’s method and Spearman distance. The optimal number of clusters (K = 5) was selected based on the consensus cumulative distribution function (CDF) plot, delta area plot, and sample tracking plot. Clustering was validated by principal component analysis (PCA), t-distributed stochastic neighbor embedding (t-SNE), and uniform manifold approximation and projection (UMAP), and silhouette analysis. For statistical analysis of cluster–category associations, a Chi-square test was performed. A p-value ≤ 0.05 was considered statistically significant.

## Results

### Establishment of the cohort and analysis of GBM infiltration by immunohistochemistry

A total of 45 IDHwt GBM patients were enrolled in the study (Figure 1A). The extent of resection was categorized according to the RANO criteria [7]: (i) supratotal resection (class 1): removal of all CE tumor and less than 5 cm^3^ of residual nCE tumor; (ii) complete resection (class 2a): removal of all CE tumor; (iii) neartotal resection (class 2b): less than 1 cm^3^ of residual CE tumor; (iv) subtotal resection (class 3a): between 1 and 5 cm^3^ of residual CE tumor; (v) partial resection (class 3b): more than 5 cm^3^ of residual CE tumor; and (vi) biopsy (class 4): no tumor reduction (Figure 1B).

**Figure 1.**
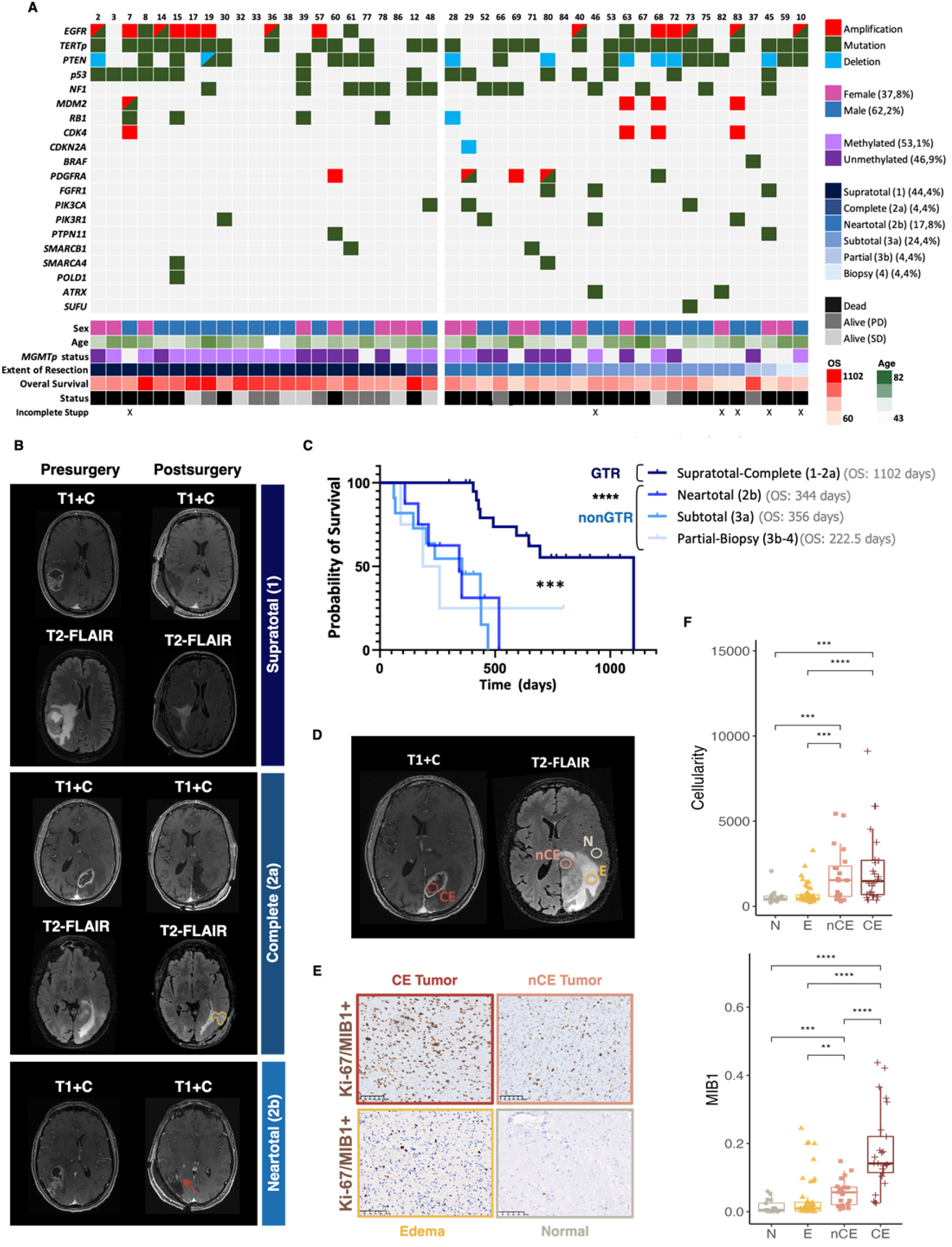
Characteristics of the patients and biopsies analyzed in this study. **A)** Oncoplot showing genetic variants in the 45 glioblastoma (GBM) tumors included in the study. Key clinical features are color-coded at the bottom. **B)** Magnetic resonance imaging (MRI) scans (T1-weighted post-contrast, T1+C; and T2 FLAIR sequence, T2+FLAIR) from three representative patients, obtained before and after surgery. The estimated extent of resection, based on postoperative MRI, is indicated on the right. In the complete resection case, residual tumor visible on T2-FLAIR is outlined in yellow. In the neartotal resection case, a red arrow highlights a residual contrast-enhancing area on T1+C. **C)** Kaplan-Meier survival curve of the patient cohort, stratified by extent of resection. **D)** Representative T1+C and T2+FLAIR images from one patient, showing potential locations of radiologically-guided biopsies (colored circles). **E)** Representative immunohistochemistry images showing MIB1 staining across different radiological regions. **F)** Quantification of cell density (top) and MIB1 labeling index (bottom) per field across the different radiological biopsy categories. PD: progressive disease, SD: stable disease. Scale bar (E): 100µm. * P≤0.05, ** P≤0.01, *** P≤0.001, **** P≤0.0001.

In our cohort, the presence of even small areas of CE tumor after surgery strongly impacted patient prognosis. No significant differences were observed between patients who underwent neartotal or subtotal surgeries (Figure 1C). To increase the statistical power of our correlation studies, we grouped patients into two categories: those who received gross total resection (GTR) of the CE tumor (supratotal and complete) and those who underwent non-GTR (neartotal, subtotal, partial, and biopsy) (Figure 1C).

We collected 161 biopsies from different radiological regions (Figure 1D). A portion of the samples was embedded in paraffin, and the number of Ki-67/MIB-1+ cells and the cellularity (total number of cells per field) were quantified automatically through IHC analysis (Figure 1E). As expected, biopsies from the CE tumor region exhibited the highest cell density and proliferation index (Figure 1F). However, nCE tumor samples showed a significantly higher percentage of MIB1+ cells than areas considered non-tumorigenic by MRI (edema and normal), and exhibited similar cellularity levels to CE biopsies (Figure 1F). Notably, among biopsies classified radiologically as edema or normal, there were instances in which the MIB1 index and/or cell density were abnormally elevated (Figure 1F).

We performed a correlation analysis between the two cellular parameters across the different radiological groups of biopsies (Figure 2A). Using the values observed in edema and normal biopsies, we defined a “pathological” range, based on the first quartile of the MIB1 index (more than 2.5%) (dotted lines in Figure 2A). As expected, all CE and most nCE samples displayed pathological features. The analysis confirmed that several biopsies classified as edema, as well as a few radiologically normal samples, displayed high MIB1 levels. Generally, signs of tumor infiltration were associated with a high MIB1 index and low cellularity in edema biopsies, and high cellularity but not as high MIB1% in nCE samples (Figure 2A). Samples with high cellularity but low MIB1% were normally categorized as reactive gliosis by pathologists (Figure 2A). Interestingly, high MIB1 levels in edema or normal samples were more prevalent among old adult GBM patients (Figure 2B), and were associated with a poorer prognosis, though this was not statistically significant (Figure 2C).

**Figure 2.**
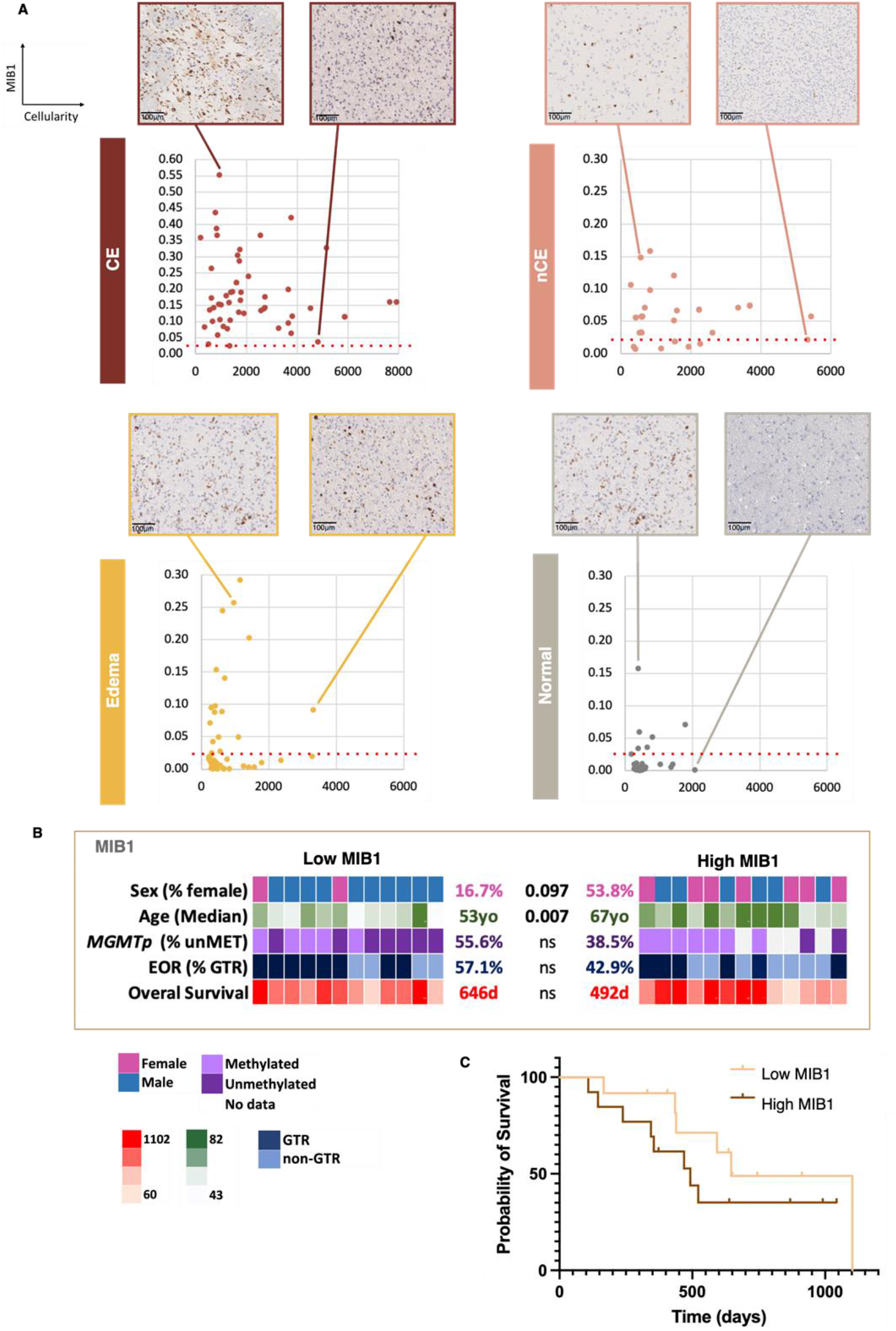
Immunohistochemical (IHC) analysis of the samples. **A)** Graphs show the MIB1 labeling index (Y-axis) and cell density (X-axis) across different radiological biopsy categories. Representative MIB1 IHC images from selected samples are displayed above the graphs. **B)** Patients were grouped based on the detection of at least one biopsy with a high MIB1 labeling index (>2.5%) in radiologically defined edema or normal regions (High MIB1 group). Patients were classified as Low MIB1 if at least two biopsies were obtained from edema or normal regions and none exceeded 2.5% MIB1+ cells. The heatmap summarizes the clinical and molecular features of the two groups. Rows represent key variables: sex, age, MGMT promoter (MGMTp) methylation status, extent of resection (EOR), and overall survival (OS). P values were calculated using Fisher’s exact test for categorical variables, the Mann Whitney test for median age, and the Log-rank (Mantel-Cox) test for OS. **C)** Kaplan–Meier survival curves comparing the High MIB1 and Low MIB1 patient groups (P=0.262B; log-rank test). ns, non-significant. Scale bar: 100µm

We analyzed the progression of some patients with signs of tumor infiltration (more than 2.5% MIB1) in biopsies that were radiologically classified as edema or normal in detail (Figure 3). Postoperative MRI revealed a small residual CE tumor area in patients PTZ28 and PTZ29 (neartotal resections) (brown arrows). However, in both cases, areas of edema with signs of tumor infiltration were also left behind after surgery (yellow arrows). Both biopsies were located close to the CE tumor: 5 mm for PTZ28 and 11 mm for PTZ29. Notably, tumor recurrence occurred very soon after surgery (six and three months, respectively) in both the residual CE tumor and the residual edema region. In contrast, patient PTZ14 underwent a supratotal resection, yet a region of edema with a low MIB1 index was left behind (yellow arrow) and was located 4 mm from the core. This patient showed signs of tumor recurrence 17.5 months after surgery in several areas far from the resection cavity and the region where the edema biopsy was taken (white arrows). These observations reinforce the notion that GBM infiltration beyond the CE and nCE regions can drive rapid tumor recurrence if not surgically removed.

**Figure 3.**
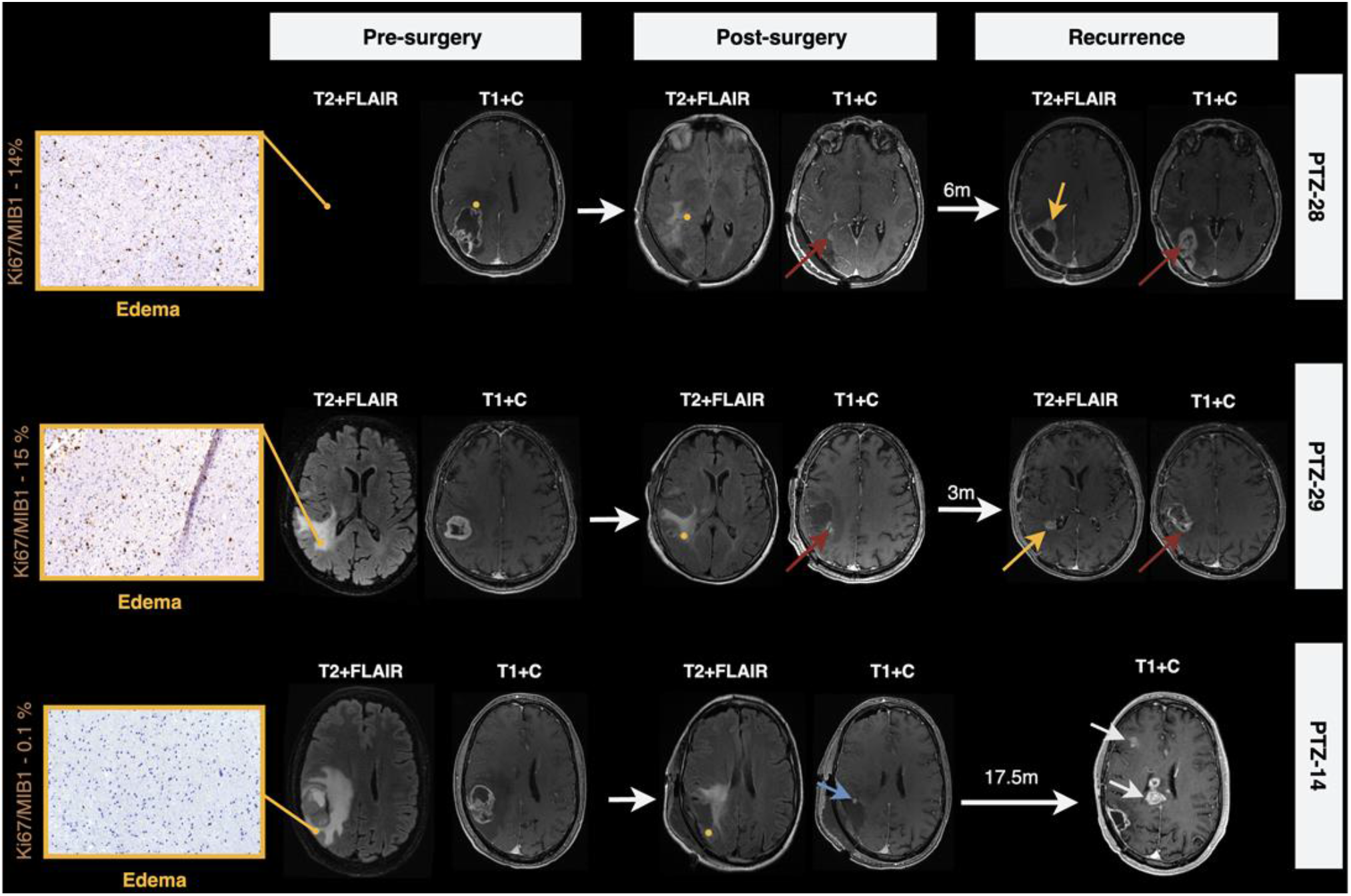
Impact of GBM infiltration into edema areas on tumor progression. Magnetic resonance images (MRI) (T1-weighted post-contrast, T1+C; and T2 FLAIR sequence, T2+FLAIR) from three representative patients are shown. Images were taken before and after surgery, as well as at the time of tumor recurrence. Yellow dots mark the locations of edema biopsies. On the left, representative MIB1 immunohistochemical staining images of these biopsies is displayed. In the post-surgical MRI, brown arrows indicate residual contrast-enhancing (CE) tumor in patients PTZ28 and PTZ29, while a blue arrow marks an ischemic region in PTZ14. In the recurrence MRI of PTZ28 and PTZ29, yellow arrows highlight new CE tumor arising from the location of infiltrated edematous tissue, and brown arrows indicate regrowth from residual CE tumor. In PTZ14, white arrows point to sites of tumor recurrence distant from the resection cavity.

### Molecular and cellular analysis of the cohort of biopsies

The tumor microenvironment (TME) at the periphery of GBM tumors differs significantly from the tumor bulk. The periphery consists primarily of normal brain tissue, while the bulk consists of hypoxic, necrotic, and angiogenic regions. This suggests that tumor cells face distinct fate-choice pressures in these two areas. To better characterize the GBM TME, we selected specific genes representative of each cell type and measured their expression in our biopsy cohort as an indication of their cellular phenotype. Using an integrated atlas from publicly available scRNA-seq databases [14] (Supplementary Figure 1), we identified distinct clusters of immune cells (myeloid and lymphoid), astrocytes, oligodendrocytes, and vascular cells (endothelial and pericytes) (Figure 4A). We also detected two clusters (glial and myeloid cells) that shared many genes overexpressed in gliomas, all of which were related to cell proliferation (Figure 4A). After several refinement steps, we selected a gene set specific to each cell type (Supplementary Figure 2) and confirmed its differential expression across distinct clusters (Figure 4B). We then measured the expression of the selected genes using qRT-PCR analysis to evaluate the distribution of individual genes and cellular scores (gene sets) in our cohort of radiologically guided GBM biopsies.

**Figure 4.**
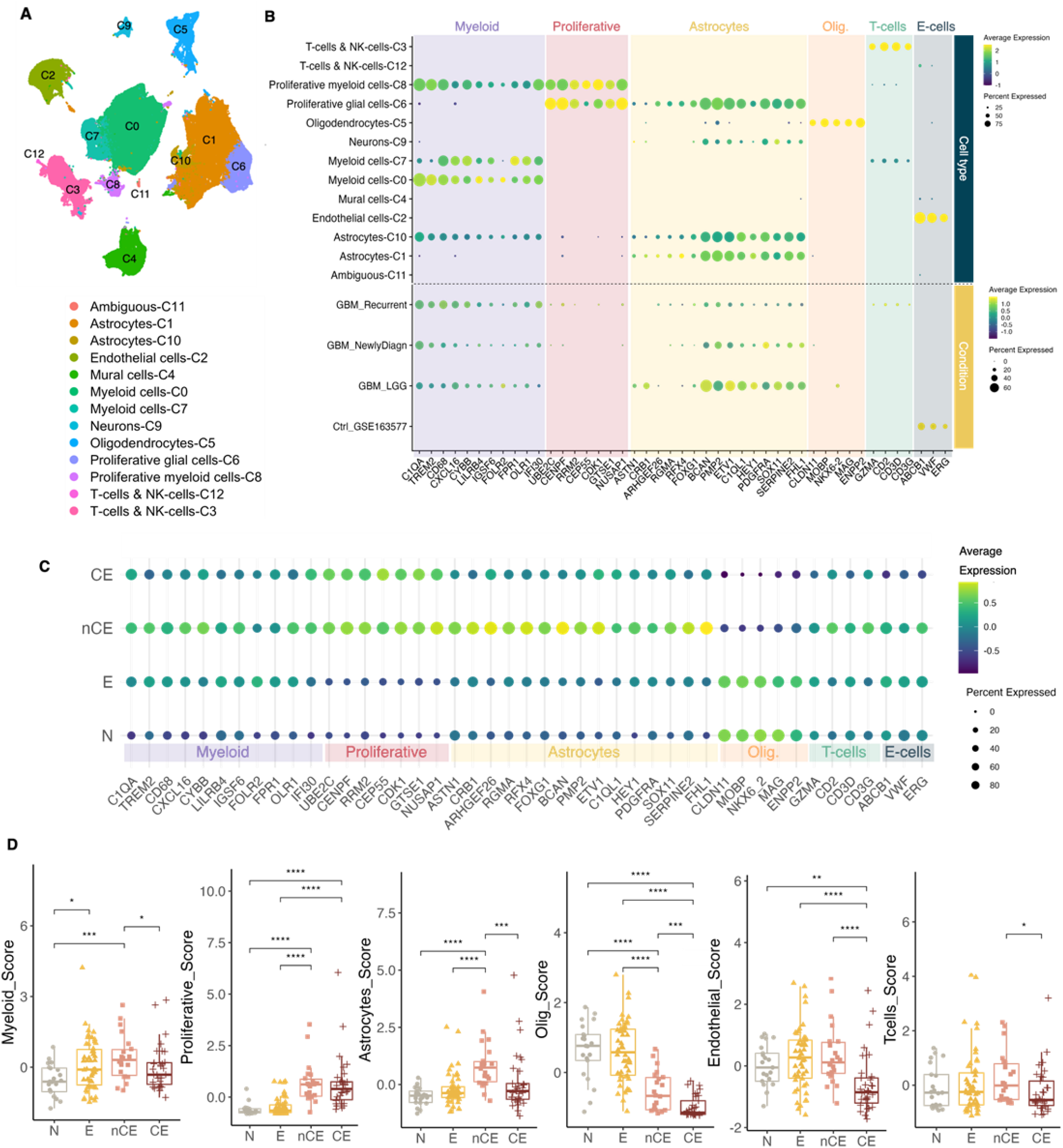
Selection of the gene panel and characterization of GBM biopsies. **A)** UMAP plot showing cell type annotation in the integrated dataset, which includes control and glioma scRNAseq data. **B)** Dot plot displaying the percentage of cells expressing each selected marker (dot size) and the average expression level (color intensity) across different cell clusters and conditions: control brain, low-grade gliomas (LGG), newly-diagnosed glioblastoma (ndGBM) and recurrent glioblastoma (rGBM). The minimum dot size threshold was set to 0.01, indicating that only genes expressed in more than 1% of cells are shown. **C)** Bar plots illustrating the log-transformed mean expression (*scores*) of each cell type across distinct radiological regions: Normal (N), Edema (E), Non-Contrast Enhancing Tumor (nCE), and Contrast-Enhancing Tumor (CE). **D)** Boxplot showing cell-type-specific scores across the different sample categories. Statistical significance was assessed using the Kruskal-Wallis test. * P≤0.05, ** P≤0.01, *** P≤0.001, **** P≤0.0001.

Consistent with the IHC findings, both CE and nCE tumor samples exhibited high expression of proliferative markers. Furthermore, a significant decrease in oligodendrocyte gene expression distinguished these two tumorigenic regions from other biopsies (Figure 4C-D). nCE tumor biopsies exhibited enrichment of astrocytic, myeloid, endothelial, and T-cell markers compared to CE samples. Additionally, myeloid-related gene expression was lower in normal biopsies than in edema samples; however, this was the only significant difference between the two regions (Figure 4C-D).

Since signs of tumor infiltration in biopsies that were radiologically classified as edema or normal were more prevalent in older adult patients, we analyzed potential age-related differences in the cellular content. However, only few genes showed a differential expression in nCE tumor biopsies, as well as in edema and normal samples in patients aged 65 and older compared to patients 64 and younger (Supplementary Figure 3A). Next, we compared samples from female and male patients and observed striking differences in the expression of several astrocytic and oligodendrocyte genes in nCE samples (Supplementary Figure 3B). Furthermore, multiple myeloid-related genes were upregulated in edema and normal biopsies (based on the radiological classification) from female GBM patients (Supplementary Figure 3B). Notably, no significant sex-related changes were observed in CE samples. These findings suggest the existence of important differences in the cellular composition of the TME outside the enhancing tumor between male and female GBM patients.

### Unsupervised clustering of the samples

Unsupervised clustering of the qRT-PCR expression data confirmed that the genes selected for the different cell types clustered together along the x-axis (Figure 5A). For sample classification, we grouped them into five clusters, which were supported by cumulative distribution function (CDF), delta area, and sample tracking plots. These plots indicated diminishing gains in cluster separation beyond K = 5 (Supplementary Figure 4A). Cluster robustness and separability were further confirmed through principal component analysis (PCA) and uniform manifold approximation and projection (UMAP), which demonstrated clear and consistent grouping of samples (Supplementary Figure 4B).

**Figure 5.**
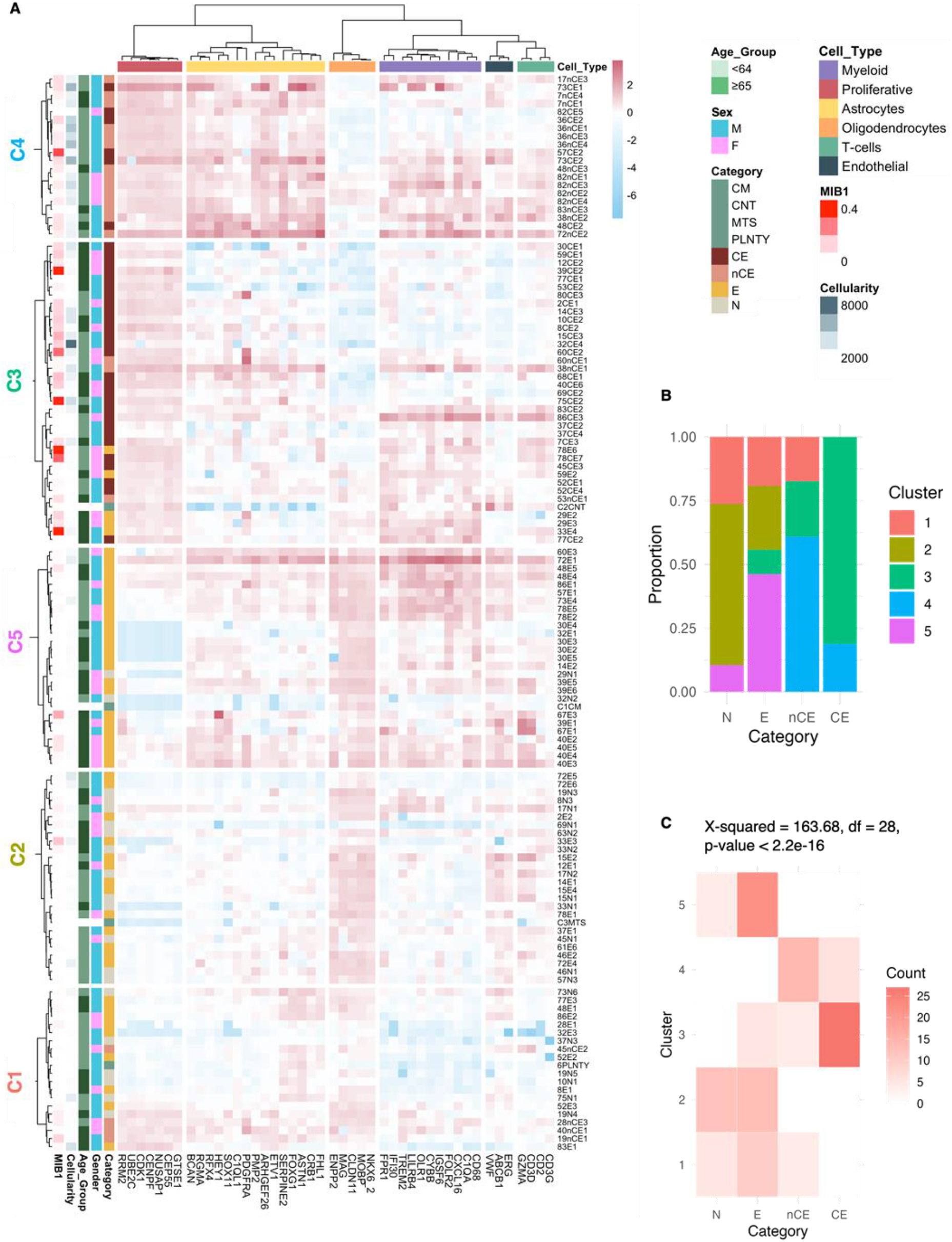
Classification of GBM biopsies using the cellular gene panel. **A)** Heatmap showing the unsupervised clustering of samples based on the expression the selected gene panel. **B)** Stacked bar plot representing the proportion of clusters (1–5) within each radiological tissue category (N: Normal, E: Edema, nCE: Non-Contrast-Enhancing Tumor, CE: Contrast-Enhancing Tumor). **C)** Heatmap illustrating the distribution of tissue categories across clusters, with color intensity indicating the number of cells per category. A chi-square test (X^2^ = 163.68, df = 28, p < 2.2e-16) confirms a significant association between clusters and tissue types.

We observed a high degree of similarity between CE and nCE tumor samples, which clustered together in Cluster 3 (C3) and Cluster 4 (C4) (Figure 5A-C). Both clusters were enriched in proliferative markers and exhibited lower expression of normal oligodendrocyte markers than the other three clusters. However, samples in C4 displayed higher levels of proliferative, astrocytic, and endothelial cell markers (Supplementary Figure 5). Notably, some samples classified radiologically as edema clustered with CE and nCE biopsies in C3. These samples exhibited high levels of MIB1% and cellularity, reinforcing the notion that these regions may represent areas of tumor infiltration (Figure 5A-C). However, most peritumoral samples were distributed into the other three groups: C1, C2, and C5. C2 showed higher levels of oligodendrocyte and T-cell markers, while C5 exhibited higher levels of myeloid markers (Supplementary Figure 5). The chi-square test confirmed a significant association between cluster assignment and sample category (X^2^ = 163.68, df = 28, p < 2.2e−16) (Fig. 5C).

To get an idea of the normality of the GBM periphery, we included a few samples from radiologically normal regions of patients with non-GBM diseases. These included one case of cavernous malformation (CM), one case of contusion (CNT), one case of brain metastasis (MTS), and one case of polymorphous low-grade neuroepithelial tumor (PLNTY). Notably, the CNT biopsy clustered together with several nCE and edema biopsies in C3, showing high levels of proliferative and myeloid markers (Figure 5A). In contrast, the other three biopsies clustered with non-tumorigenic biopsies in C5 (CM), C2 (MTS), and C1 (PLNTY). These biopsies showed low proliferative and high endothelial cell expression (Figure 5A).

### Prognosis value of the expression of the gene panel

To further investigate the clinical relevance of the gene panel further, we performed a correlation analysis of each gene’s expression level and disease progression. We also stratified patients into two groups based on the extent of resection. We focused on edema and CE samples because these groups had the largest sample sizes. In CE biopsies, we found that many immune, vascular, and oligodendrocyte genes were negatively correlated with prognosis in patients who underwent GTR (Figure 6A). These results suggest that the tumor core phenotype impacts disease progression, even after a complete CE resection. Conversely, higher expression of certain immune-related and proliferative genes was associated with improved survival in patients who underwent non-GTR (Figure 6A). These results imply that, in the presence of residual CE disease, these markers may be associated with a better response to radiotherapy and/or temozolomide.

**Figure 6.**
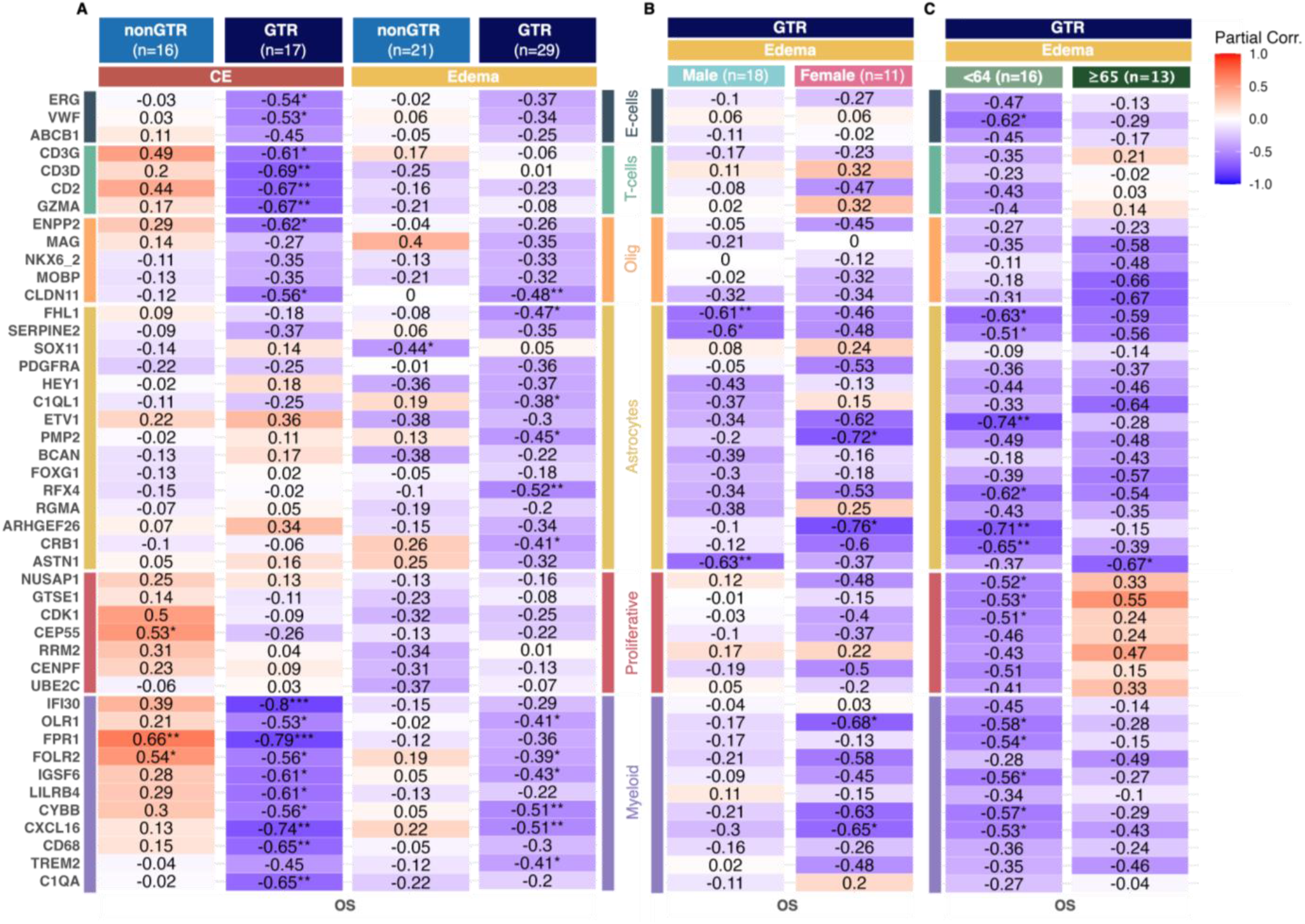
Association of specific gene markers with patient survival. **A)** Heatmap showing the correlation between the expression of selected genes in contrast-enhancing (CE) tumor and edema regions with overall survival. Patients were stratified by extent of resection: Gross Total Resection (GTR) vs non-GTR. Genes are grouped by associated cell types: endothelial cells (E-cells), T-cells, oligodendrocyte cells (Olig.), astrocytes, proliferative cells, and myeloid cells. **B)** Heatmap displaying the correlation between gene expression in edema regions and overall survival in male and female GBM patients (left panels) and younger and older GBM patients (right panel) who underwent GTR. The survival analysis was adjusted for censoring. * P≤0.05, ** P≤0.01, *** P≤0.001.

The expression of many genes, including myeloid, astrocyte, and oligodendrocyte markers, in edema biopsies also negatively correlated with disease progression, particularly among GTR patients (Figure 6A).

Considering the previously observed differences in the cellular phenotype of male and female patients, as well as the distinct level of tumor infiltration in the periphery in younger versus older patients, we analyzed the impact of edema gene expression separately in the GTR patient subgroups. Notably, a significant inverse correlation of the myeloid expression with survival was observed only in female patients, whereas higher expression of astrocytic markers correlated with worse survival in all GBM patients (Figure 6B). Regarding age, the most striking difference was the positive correlation (though not statistically significant) of proliferative gene expression in the edema with OS among older GBM patients. In patients age 64 and younger, the expression of many myeloid, proliferative, astrocytic and endothelial cells, was significantly associated with poorer prognosis.

Taken together, these findings suggest that the nature of the TME and the degree of tumor infiltration in the peripheral zone of GBM have distinct effects on disease progression. These effects vary depending on the patient’s sex and age, as well as the extent of resection.

## Discussion

GBM remains one of the most challenging cancers to treat, with only a few successful clinical trials conducted in recent decades. One approach that has significantly improved OS in patients with GBM, is extending surgical resection to include nCE tumor regions [5-7]. Our findings reinforce the value of such supratotal resections because nCE tumor biopsies exhibit cellular characteristics, particularly with regard to cell density and proliferative gene expression, that are comparable to those found in CE regions. However, MIB1 staining is lower in nCE regions, while glial markers are elevated compared to CE tumor regions. This suggests distinct tumor phenotypes in enhancing and non-enhancing compartments [18, 19]. Notably, tumor cells in nCE regions have previously been shown to express LGG-associated genes [20], and in our cohort, selected astrocyte genes, with high expression levels in LGG, were also highly expressed in nCE biopsies.

Due to the small size of our study group and the fact that many patients were still alive at the time of the analysis, we could not definitely confirm the benefits of supratotal resection. Nevertheless, we observed a clear survival difference between patients who underwent GTR and those with less than 1 cm^3^ of CE tumor post-surgery. A prior Norwegian study previously documented a significant survival benefit of complete CE resection compared to neartotal resection [21], suggesting that even minimal residual CE tumor may affect disease progression and/or treatment response. Further studies in larger cohorts are needed to validate these findings.

A major challenge in performing more extensive GBM resections is distinguishing nCE tumor regions from edema because both appear as hyperintense regions on FLAIR sequences. This distinction often relies on subjective radiological criteria and becomes particularly difficult after surgery [7, 22]. Importantly, our data indicate that areas previously considered to be vasogenic edema, or even normal tissue, on MRI may in fact harbor significant tumor infiltration. These findings underscore the urgent need for improved intraoperative tools to delineate infiltrated peripheral regions of GBM more effectively. However, our results also suggest exercising caution when applying the concept of supratotal resections to elderly patients because infiltrated tumor tissue hidden within high FLAIR signal regions was observed more frequently in older individuals. This finding may help explain why extended resections do not appear to confer clinical benefit in patients over 65 years of age [5]. It is also consistent with recent studies indicating that molecular and cellular alterations in the extratumoral zone, rather than in the CE tumor core, may better account for the poorer prognosis observed in elderly GBM patients [23, 24]

The correlation between the MIB1 labeling index and cellularity values suggests a differential balance between proliferation and migration across distinct areas of GBM infiltration. Infiltrated edema biopsies exhibit particularly high MIB1 indices and low cellularity. Recent findings have shown that migratory GBM cells retain a high proliferative capacity, resulting in regions characterized by elevated MIB1 expression but reduced cellularity [25]. Our results support the notion that distinct regulatory mechanisms may govern tumor behavior depending on the anatomical location. However, we cannot rule out the contribution of non-proliferative stromal cells (e.g., reactive astrocytes or microglia), which may accumulate around GBM cells in nCE tumor regions and increase the observed cell density in these areas.

Much GBM research has focused on the core of the tumor, where growth mechanisms are heavily influenced by microenvironmental pressures, such as hypoxia, necrosis, and angiogenesis. However, this focus may be less relevant for patients who undergo GTR. In these cases, recurrence is primarily driven by residual tumor cells infiltrating the surrounding brain parenchyma. These cells are not subjected to the same selective pressures as the core. Despite the theoretical importance of this distinction, clinical trials evaluating treatment responses based on residual tumor volume remain scarce. A seminal study by Ellingson et al. identified postoperative residual CE tumor volume as an independent prognostic factor for OS in patients with newly diagnosed GBM, irrespective of therapy type [26]. Notably, long-term survivors were disproportionately represented among patients with no residual CE tumor treated with vorinostat, a histone deacetylase inhibitor [26, 27]. Conversely, some immunotherapies have shown greater efficacy in patients with residual CE tumor after surgery [28]. In line with these observations, we found that the expression of immune-related genes in the tumor core was positively associated with OS in patients with residual CE disease, but negatively associated in patients who underwent GTR. These findings suggest that the therapeutic vulnerabilities of core and infiltrative tumor cells may differ substantially, a concept supported by previous studies [29]. These results highlight the importance of stratifying GBM patients based on the presence or absence of residual CE tumor and tailoring treatment strategies accordingly, considering them as distinct clinical entities.

Regarding sex-based differences, although the higher incidence of GBM in males is well-established, less is known about brain-wide and systemic sex differences in tumor biology or treatment response. For instance, sexual dimorphism has been observed in the function of macrophages in both traumatic brain injury [30] and GBM [31, 32]. Furthermore, recent studies suggest that male and female GBM patients may experience survival outcomes governed by different biological mechanisms [33]. The differences found in the composition of the TME and the correlation analyses align with this notion. In female patients, myeloid and astrocytic gene expression in edema biopsies negatively correlated with OS, whereas in male patients, only astrocytic gene expression appeared to have a significant impact. These findings support the idea that GBM induces both micro- and macroenvironmental changes in the brain and that such extrinsic alterations, as well as intrinsic factors such as GBM stem cell phenotypes, may be modulated by sex.

Many factors have been suggested to explain the high failure rate of GBM clinical trials. One factor is overreliance on surgically resected CE tissue, which may not accurately represent the tumor’s full biological heterogeneity. We hope that our study contributes to closing this gap by providing a framework for classifying the tumor margin’s diverse cellular composition. Nonetheless, we acknowledge several limitations. The relatively small cohort size may restrict the prognostic power of some molecular and cellular markers. MGMT promoter methylation status could not be assessed in all patients. Additionally, our selected gene panel cannot discriminate between functionally distinct astrocyte or myeloid subpopulations (e.g., microglia versus blood-derived macrophages or immunosuppressive versus inflammatory phenotypes), which are important in GBM progression and may exhibit spatial variability [12]. Further research is essential to refine the cellular landscape of the peritumoral zone across different GBM patient groups and ultimately aid in the discovery of novel biomarkers and therapeutic targets within the infiltrative tumor ecosystem.

## Ethics

The study was carried out in accordance with the Declaration of Helsinki and received approval from the local ethics committee (CEI_21/480). All the participating patients provided written informed consent.

## Authorship

A.P.N. and P.S.G conceived and planned the experiments; A.P.N., J.M.S.S. and P.S.G. got funding; A.R., J.M.S.S., R.G., B.S.C., G.V., P.G., L.J.R., G.G.P. provided resources and clinical data; O.D., J.R., M.A.R.G, B.H., A.A., B.M. and A.H.L. carried out the experiments; O.D., J.R., A.P.N. and P.S.G. analyzed the data and contributed to the interpretation of the results; O.D. and P.S.G. took the lead in writing the manuscript. All authors provided critical feedback and contributed to the final document.

## Data availability

The data that support the findings of this study and the materials used are available on request from the corresponding authors.

## Acknowledgements

We sincerely thank the patients and their families to their support of this project. We are also grateful to the Neurosurgery and Pathology Departments at Hospital 12 de Octubre for their assistance with the studies presented in this manuscript. Additionally, we thank Federico Gaiti, Yiyan Wu, and Derek Wainwright for their critical review of the manuscript.

