## Supplementary Figures for "Patient-Derived Surgical samples reveal patterns of glioblastoma infiltration and tumor microenvironment at the tumor margin"

A

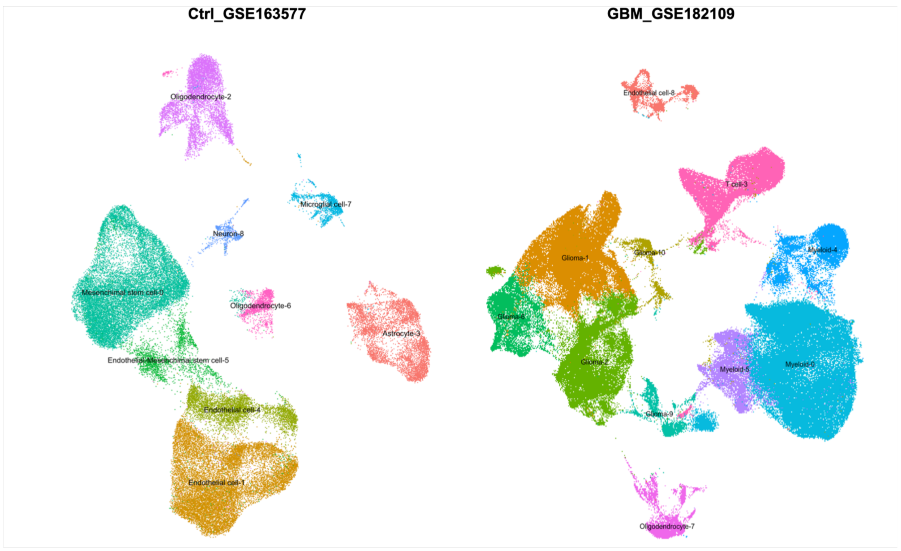


B

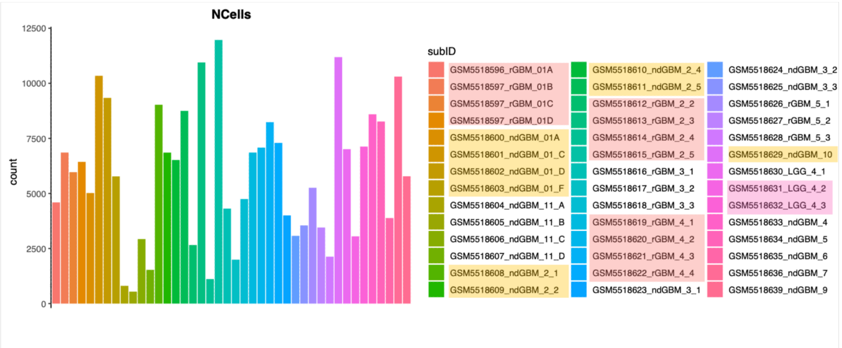


| ID | Freq |
|---|---|
| LGG_3 | 8171 |
| LGG_4 | 12940 |
| ndGBM_01 | 28722 |
| ndGBM_10 | 10655 |
| ndGBM_11 | 4339 |
| ndGBM_2 | 27580 |
| ndGBM_3 | 6370 |
| ndGBM_4 | 7989 |
| ndGBM_5 | 7256 |

| ID | Freq |
|---|---|
| ndGBM_6 | 3415 |
| ndGBM_7 | 9641 |
| ndGBM_8 | 5942 |
| ndGBM_9 | 5324 |
| rGBM_01 | 27487 |
| rGBM_2 | 23648 |
| rGBM_3 | 9908 |
| rGBM_4 | 28157 |
| rGBM_5 | 10261 |

C

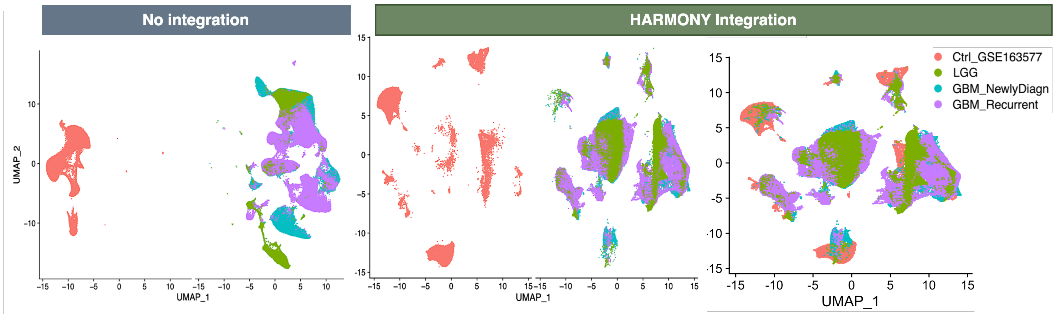


**Supplementary Figure 1. Overview of single-cell RNA sequencing (scRNA-seq) datasets and their integration. A)** UMAP projections showing the two public datasets used to construct the integrative scRNA-seq atlas. **B)** Bar plot displaying the number of cells per sample, accompanied by a color-coded sample identifier table. Below, frequency tables summarize the distribution of cells across different sample groups, including Lower-Grade Glioma (LGG), Newly Diagnosed GBM (ndGBM), and Recurrent GBM (rGBM). Highlighted samples indicate those selected for the final scRNA-seq atlas. **C)** UMAP projections comparing cell distributions before and after Harmony integration. Colors represent different datasets, including Control (Ctrl), LGG, Newly Diagnosed GBM, and Recurrent GBM

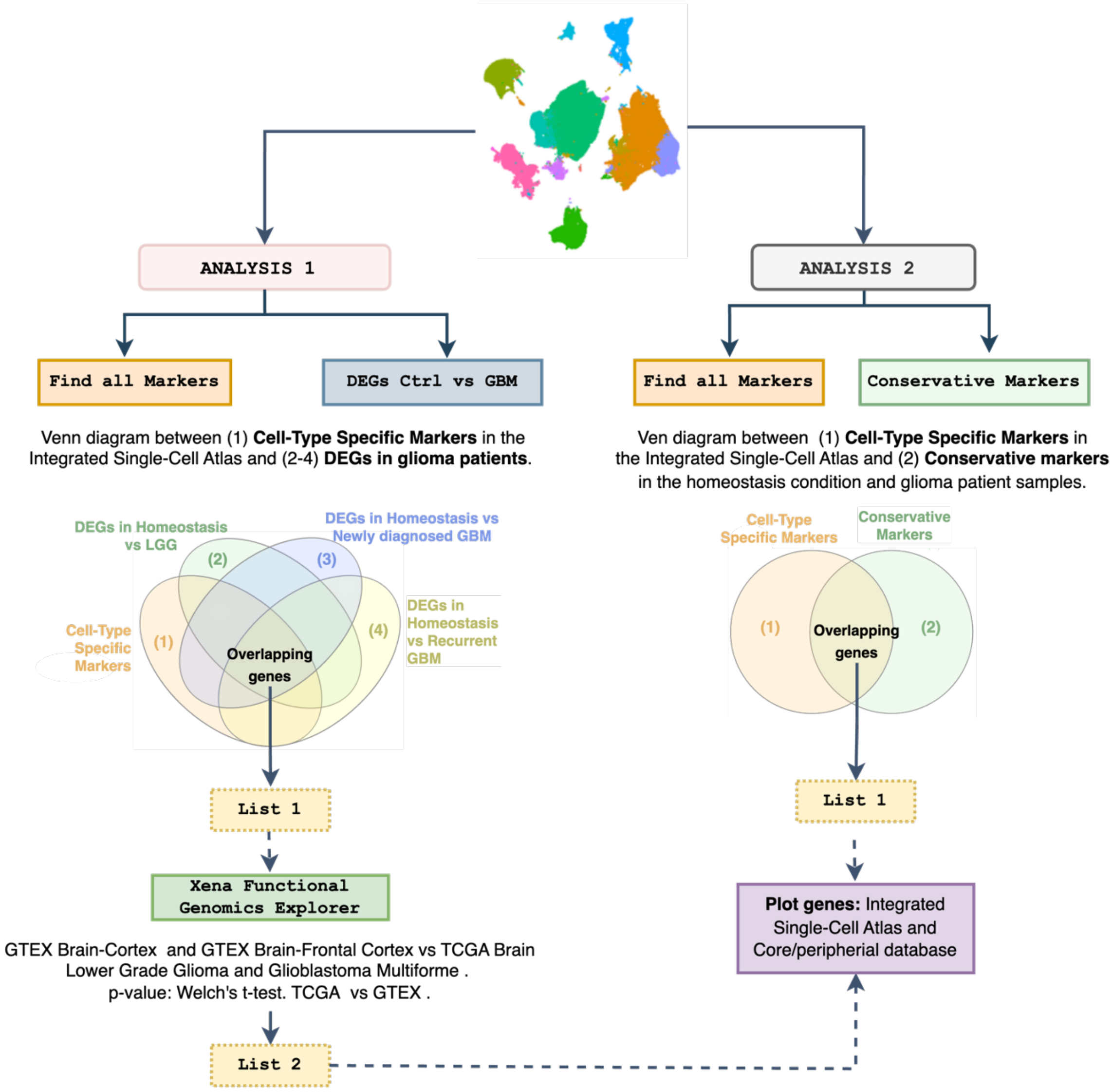


**Supplementary Figure 2. Schematic representation of the analytical workflow for identifying cell type specific gene markers in GBM using single-cell transcriptomics**. The workflow is divided into two main analyses: Analysis 1 (left) identifies differentially expressed genes (DEGs) by comparing control (Ctrl) vs. GBM and categorizing them across different conditions (homeostasis vs. lower-grade glioma (LGG), newly diagnosed GBM, and recurrent GBM). Overlapping genes are selected and further analyzed using the Xena Functional Genomics Explorer to refine List 2. Analysis 2 (right) identifies conservative and cell-type-specific markers and extracts overlapping genes. Genes from both analyses were plotted in the Integrated Single-Cell Atlas and the Core/Peripheral Database to visualize their cell type specify.

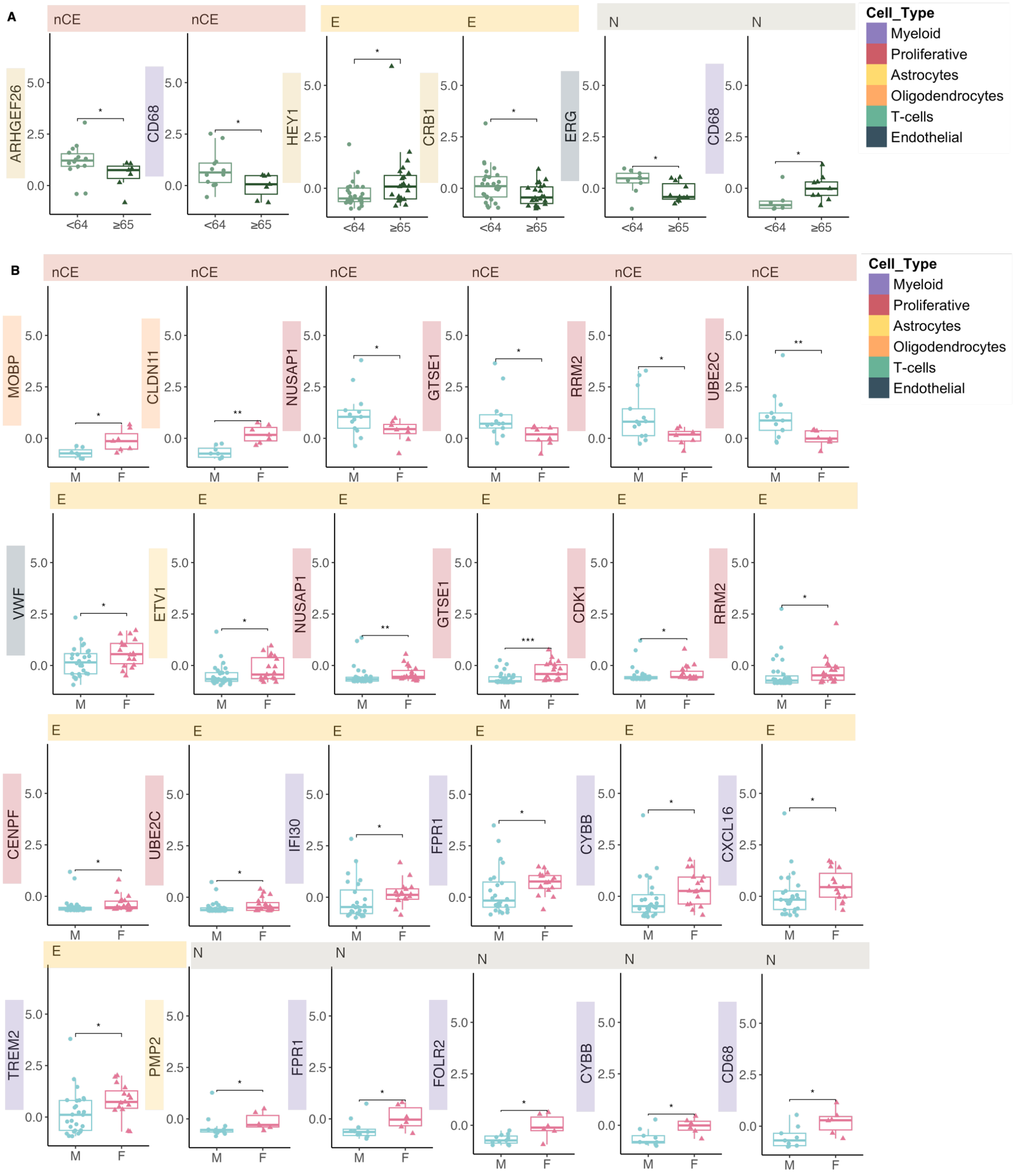


**Supplementary Figure 3. Association of specific markers with gender and age. A)** Box plots displaying the significant cell type markers between male (M) and female (F) patients, across different tumor areas: Contrast-Enhancing Tumor (CE), Non-Contrast Enhancing Tumor (nCE), Edema, and Normal tissue. Statistical significance was assessed using the Wilcoxon test. **(B)** Box plot showing the significant cell type markers differences between younger (<64 years) and older (≥65 years) patients, across the different tumor areas (CE, nCE, Edema, and Normal tissue), assessed using the Wilcoxon test. * P≤0.05, ** P≤0.01, *** P≤0.001.

A

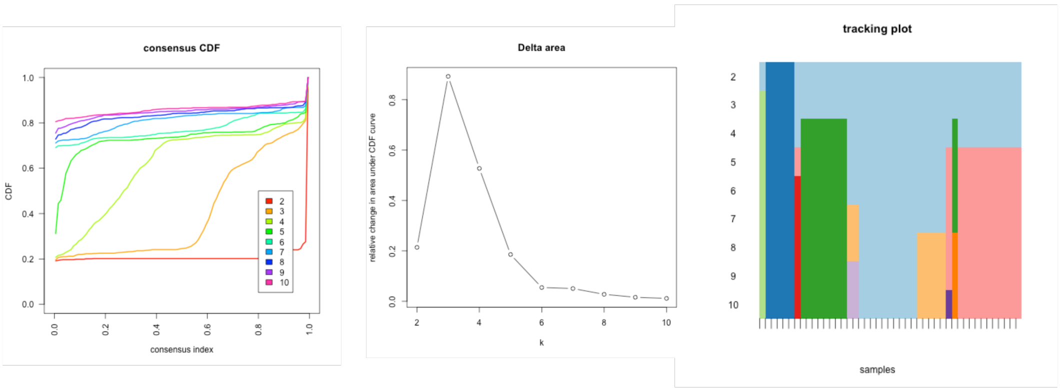


B

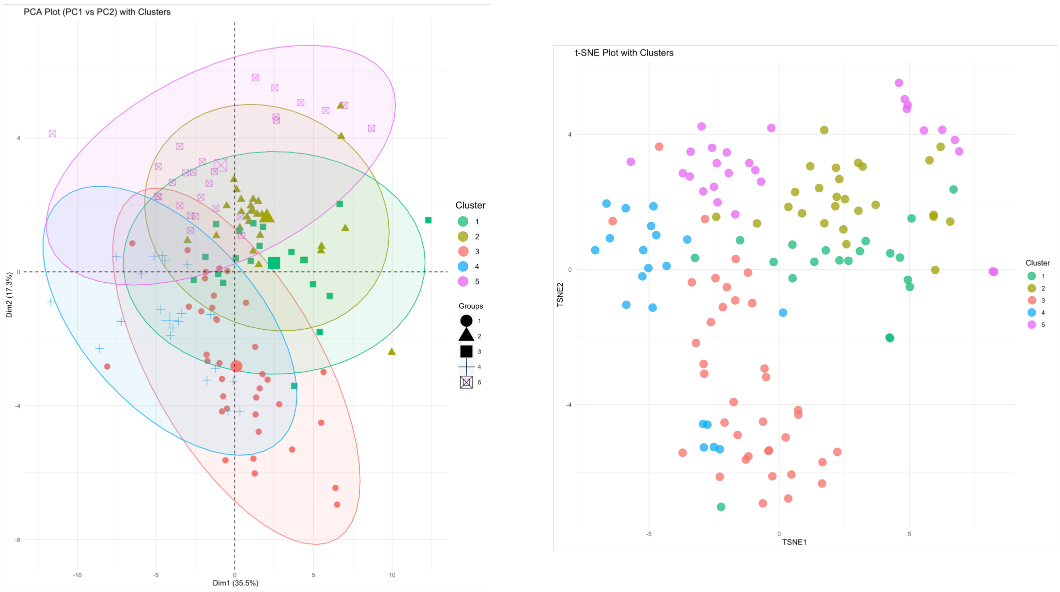


**Supplementary Figure 4. Clustering analysis and dimensionality reduction of qPCR data**. **A)** Selection of the optimal number of clusters. Consensus cumulative distribution function (CDF) plot displaying the stability of clustering solutions across different cluster numbers. Delta area plot showing the relative change in area under the CDF curve. Tracking plot illustrating the assignment of samples to clusters across different clustering solutions. **B)** Dimensionality reduction and clustering visualization. Principal Component Analysis (PCA) plot (PC1 vs. PC2) with color-coded clusters and confidence ellipses, visualizing sample distribution in reduced-dimensional space. UMAP plot showing the clustering of samples based on qPCR data, with colors representing distinct clusters.

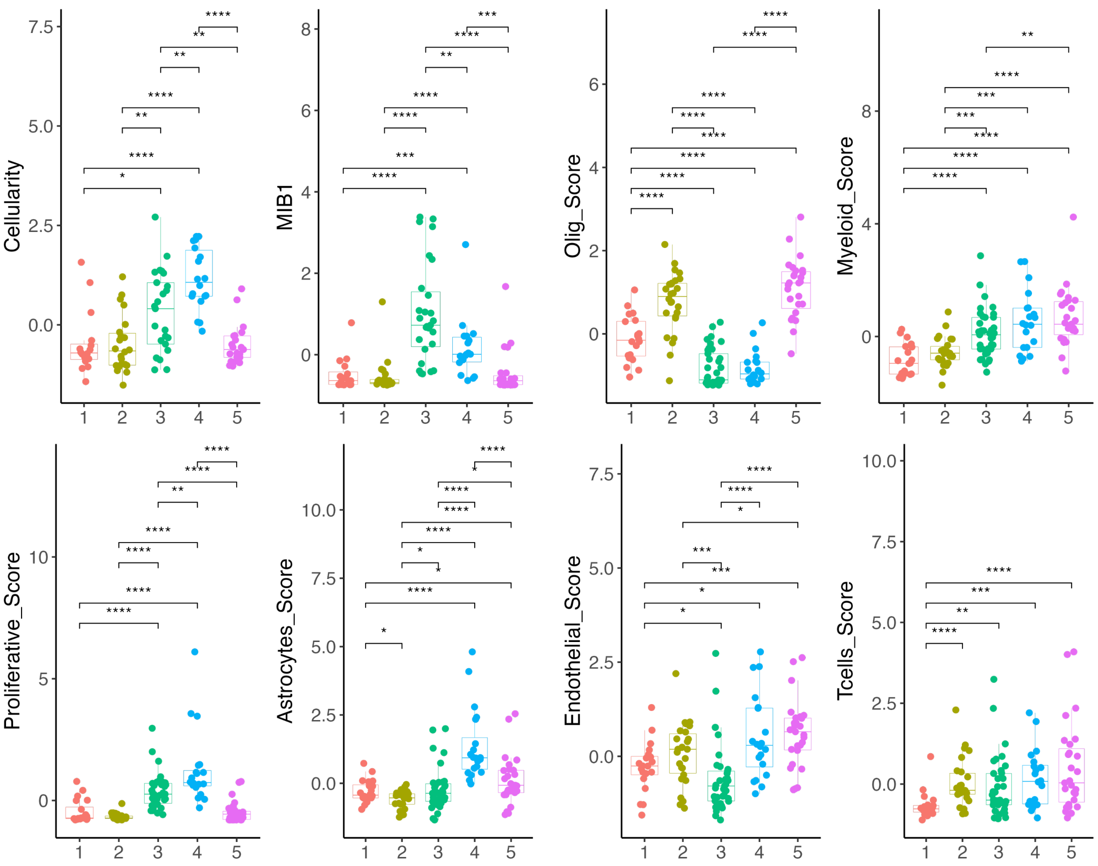


**Supplementary Figure 5. Characterization of the five identified cluster. A)** Box plots showing the mean expression of cellularity, MIB1, and cell type scores across different clusters. Statistical significance was assessed using the Kruskal-Wallis test. * P≤0.05, ** P≤0.01, *** P≤0.001, **** P≤0.0001.
