## Supplementary Data 3 for "Patient-Derived Surgical samples reveal patterns of glioblastoma infiltration and tumor microenvironment at the tumor margin"

### Analysis 1

DEGs by Condition & Find All Markers

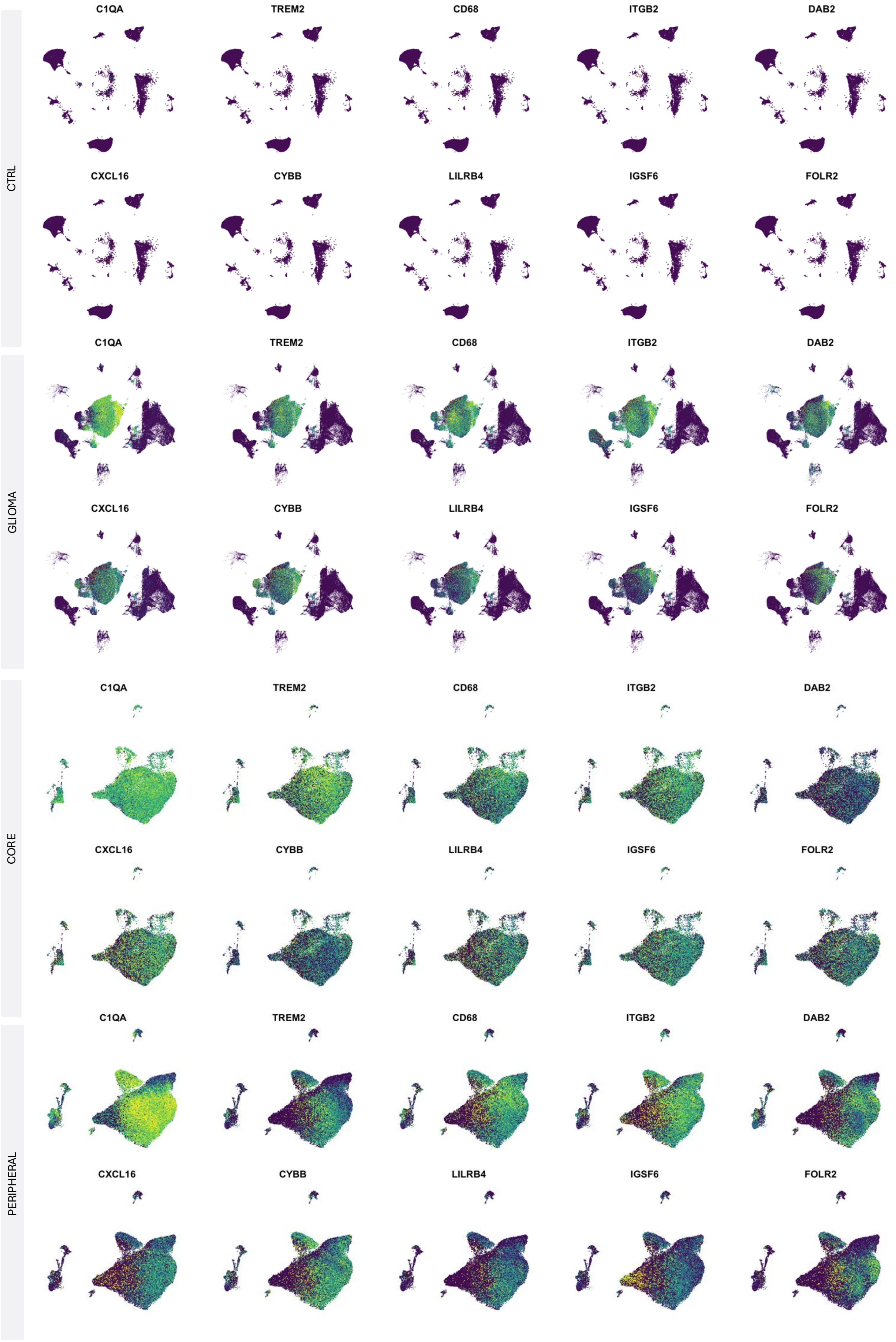
CTRL
C1QA
TREM2
CD68
ITGB2
DAB2
CXCL16
CYBB
LILRB4
IGSF6
FOLR2
GLIOMA
C1QA
TREM2
CD68
ITGB2
DAB2
CXCL16
CYBB
LILRB4
IGSF6
FOLR2
CORE
C1QA
TREM2
CD68
ITGB2
DAB2
CXCL16
CYBB
LILRB4
IGSF6
FOLR2
PERIPHERAL
C1QA
TREM2
CD68
ITGB2
DAB2
CXCL16
CYBB
LILRB4
IGSF6
FOLR2

### C1-Glioma/Astrocyte

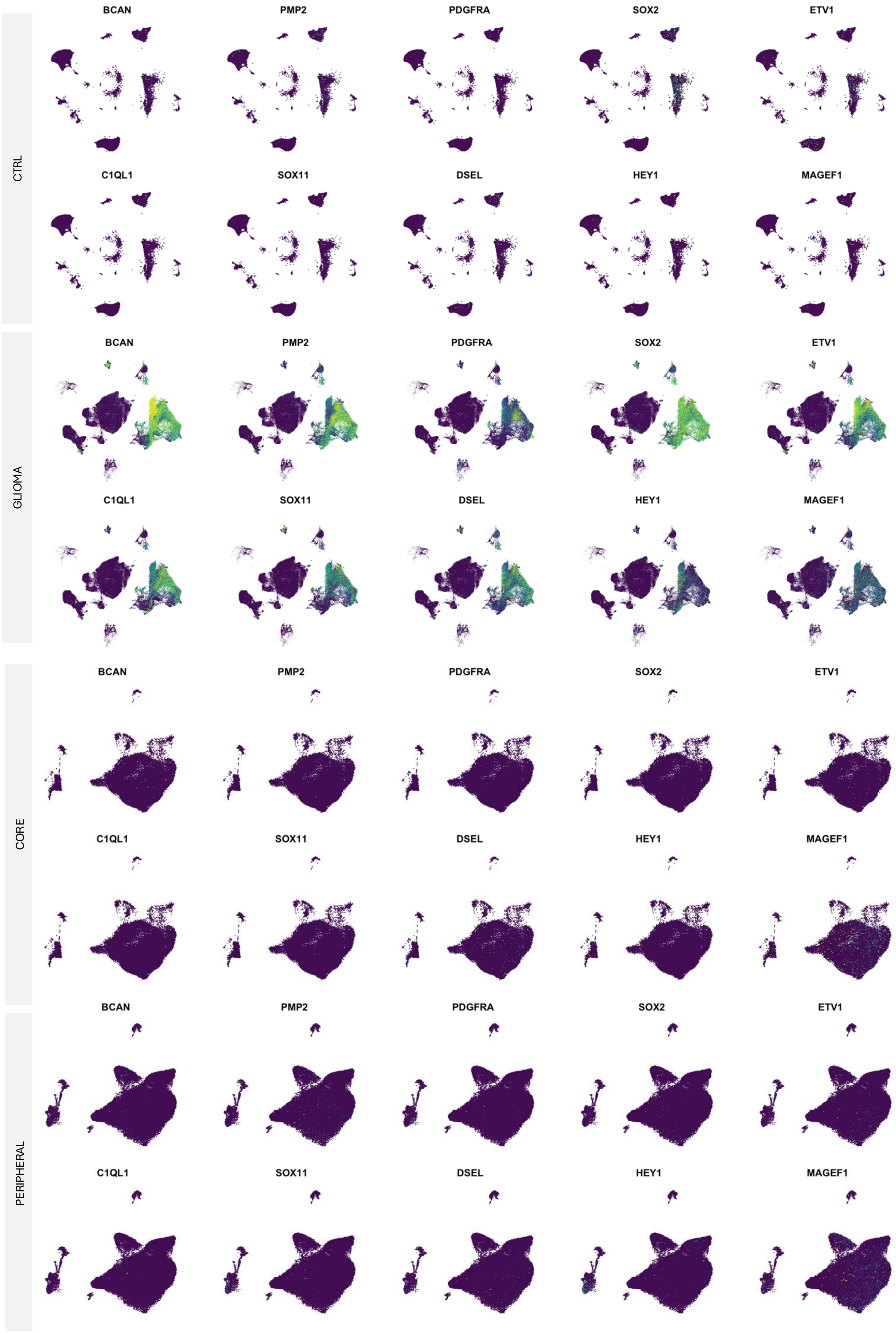

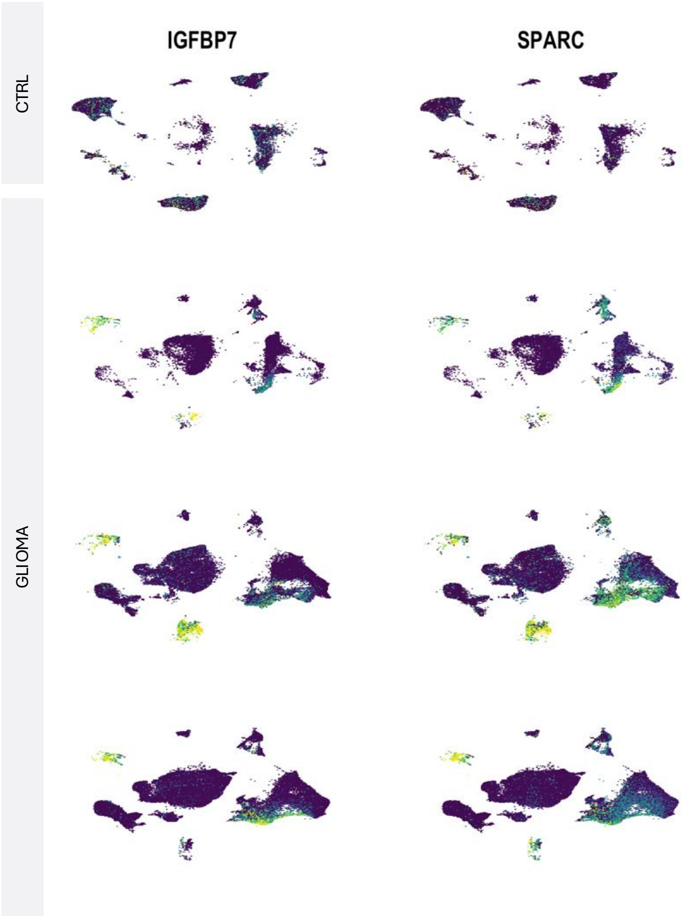
IGFBP7
SPARC
CTRL
GLIOMA

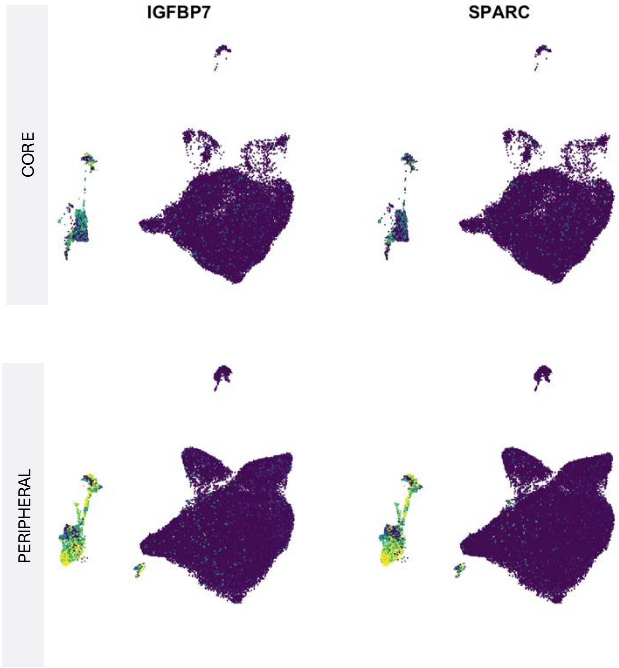
IGFBP7
SPARC
CORE
PERIPHERAL

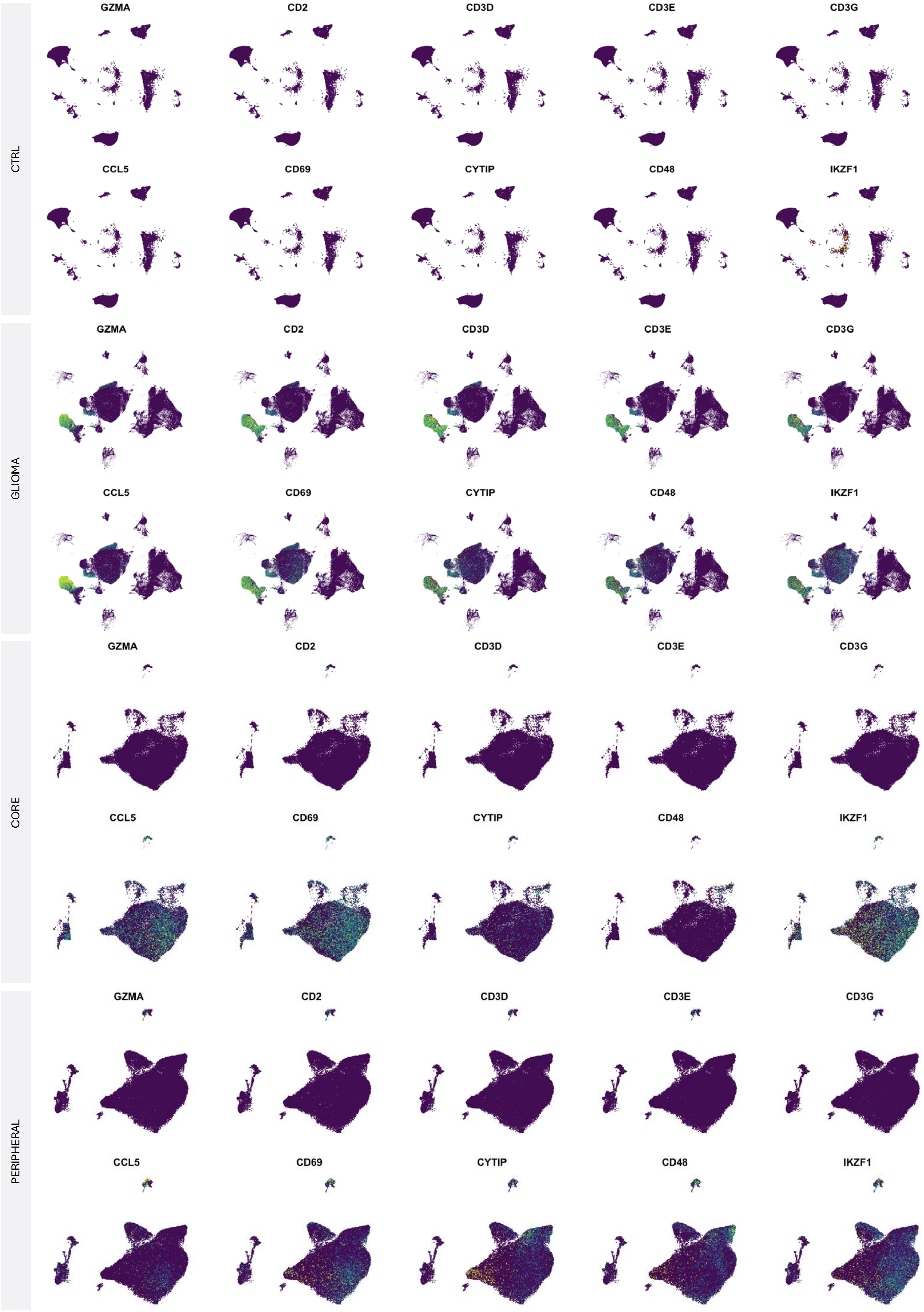
CTRL
GZMA
CD2
CD3D
CD3E
CD3G
CCL5
CD69
CYTIP
CD48
IKZF1
GLIOMA
GZMA
CD2
CD3D
CD3E
CD3G
CCL5
CD69
CYTIP
CD48
IKZF1
CORE
GZMA
CD2
CD3D
CD3E
CD3G
CCL5
CD69
CYTIP
CD48
IKZF1
PERIPHERAL
GZMA
CD2
CD3D
CD3E
CD3G
CCL5
CD69
CYTIP
CD48
IKZF1

#### C4-Mural cell

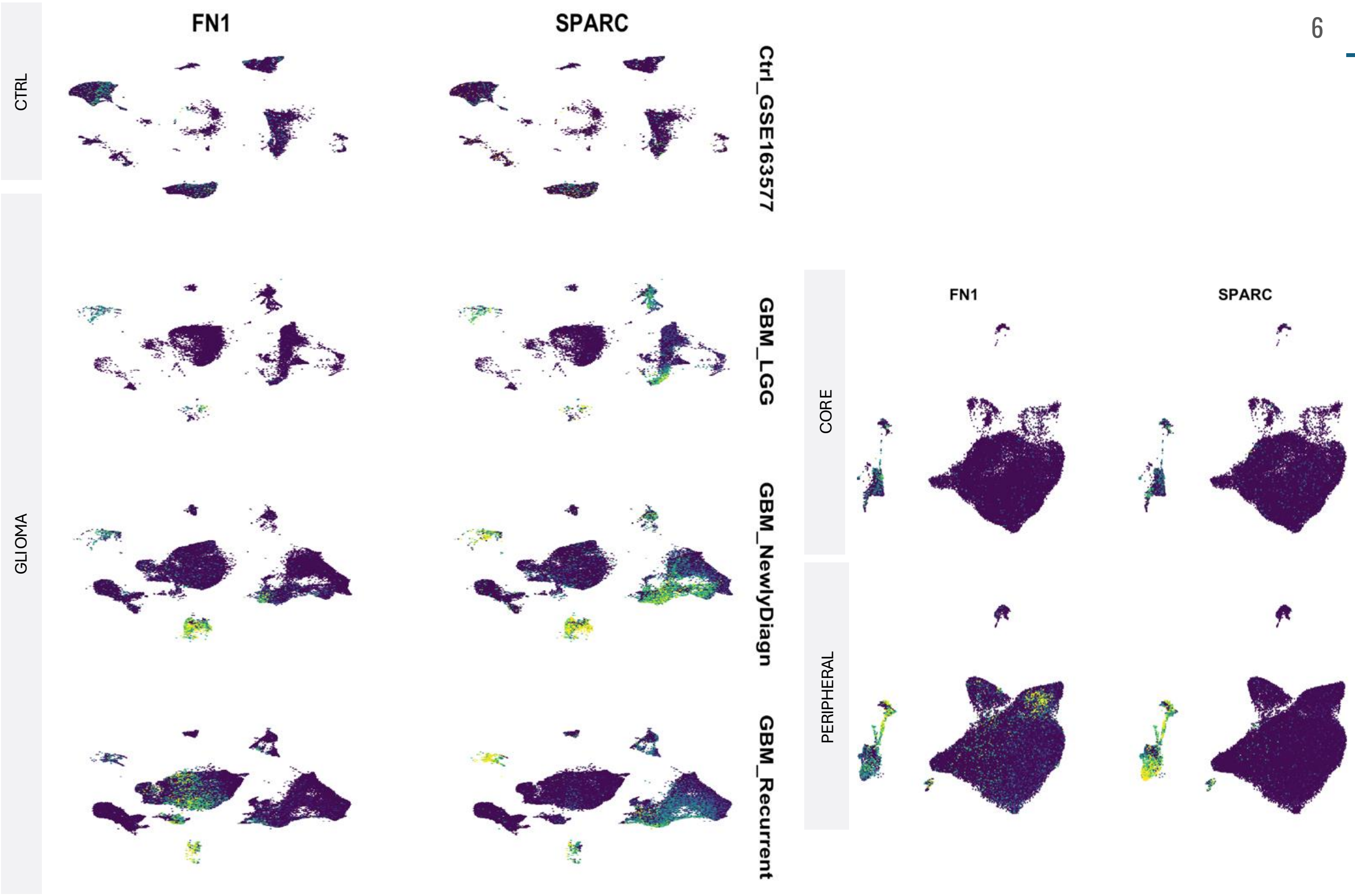


#### C5-Oligodendrocyte

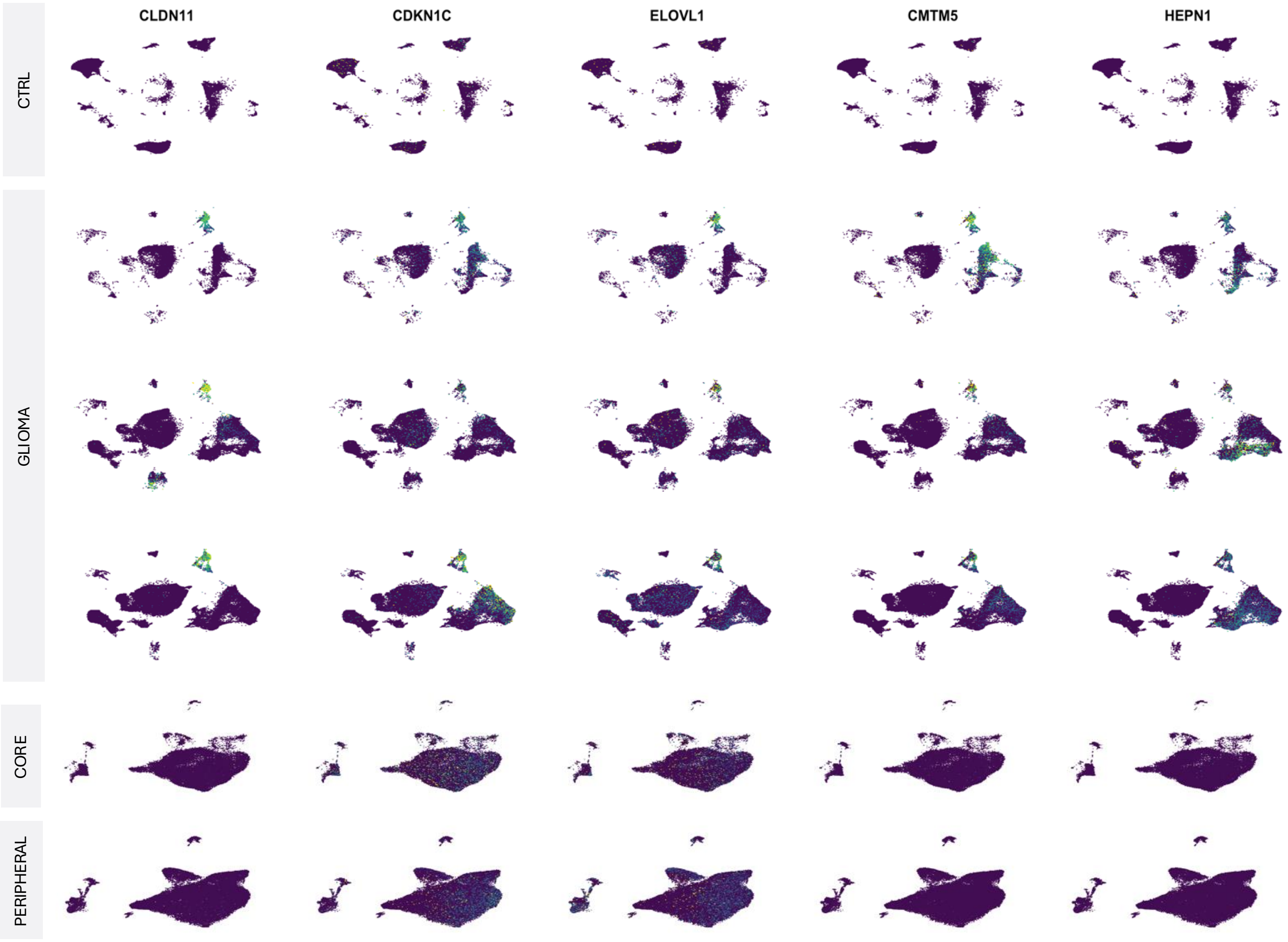

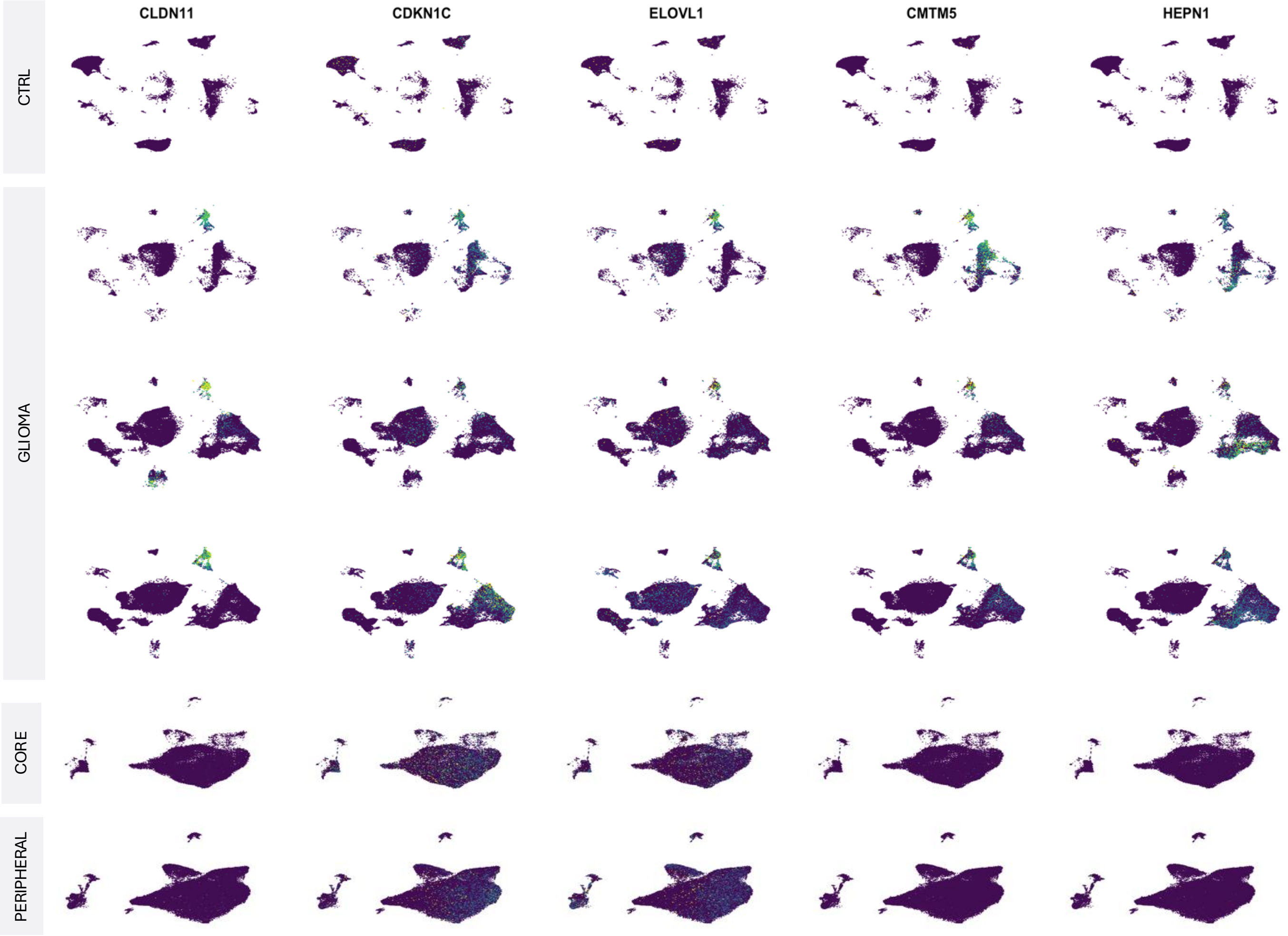
CLDN11
CDKN1C
ELOVL1
CMTM5
HEPN1
CTRL
GLIOMA
CORE
PERIPHERAL

C6- Proliferative Glial

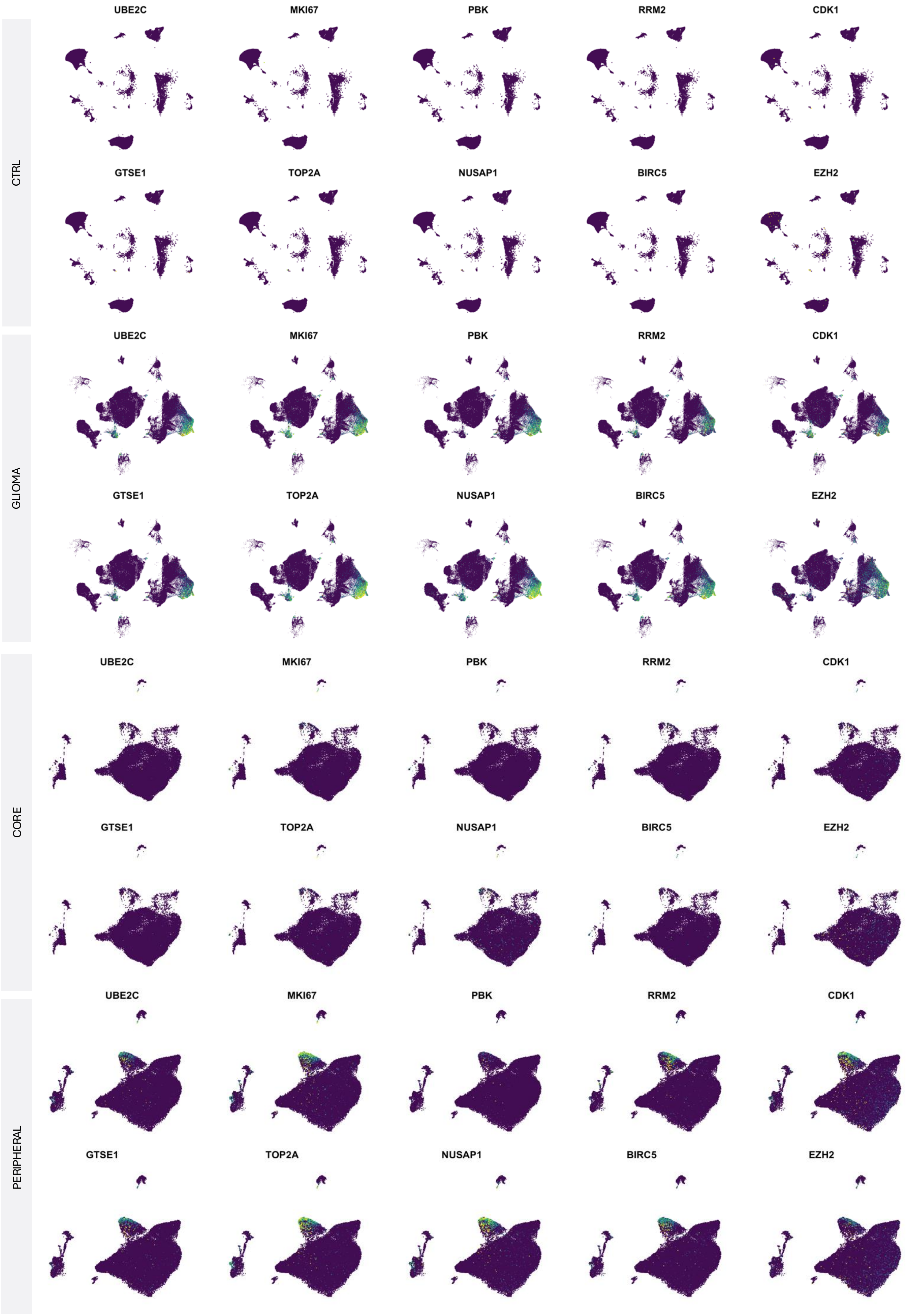

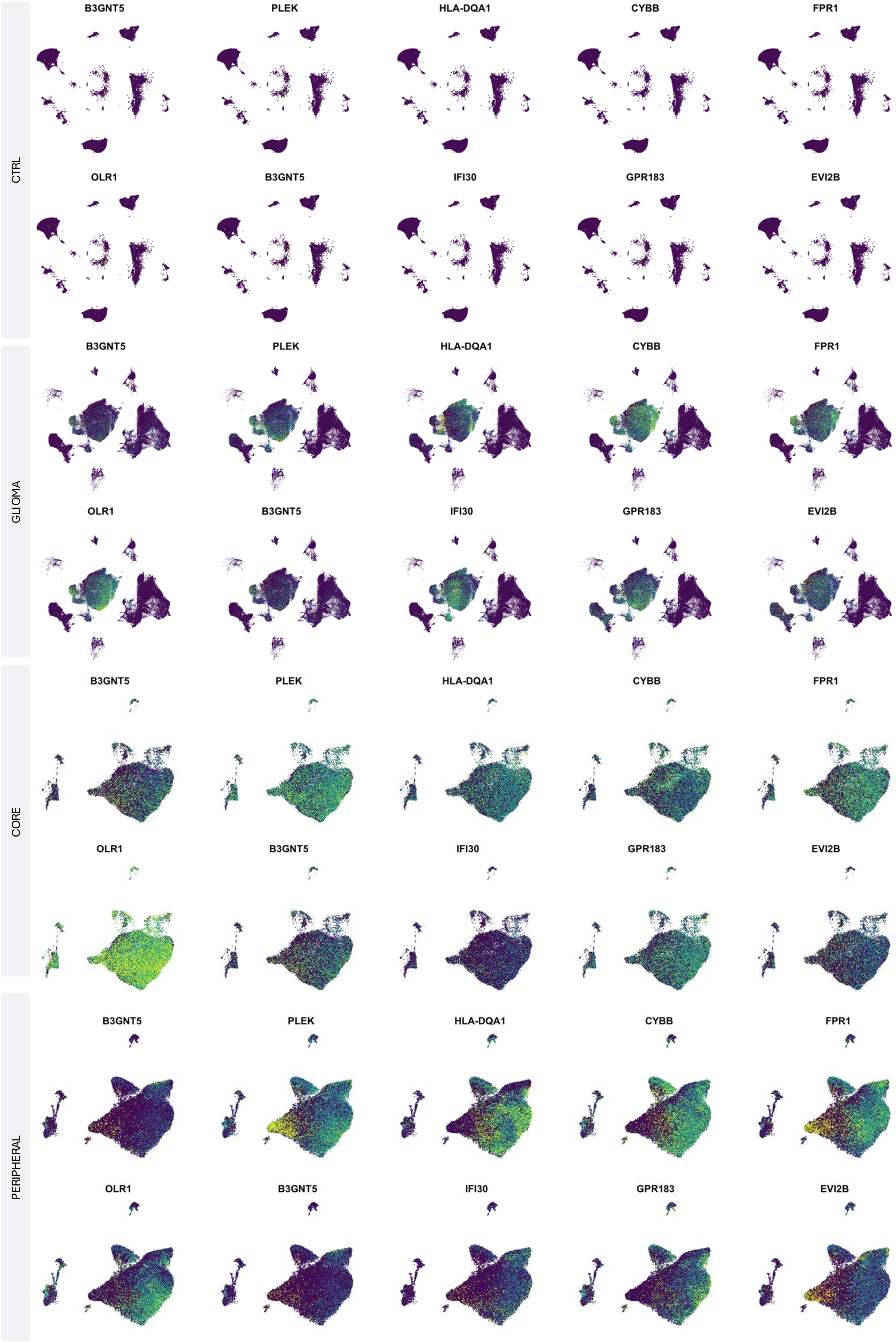
CTRL
B3GNT5
PLEK
HLA-DQA1
CYBB
FPR1
OLR1
B3GNT5
IFI30
GPR183
EVI2B
GLIOMA
B3GNT5
PLEK
HLA-DQA1
CYBB
FPR1
OLR1
B3GNT5
IFI30
GPR183
EVI2B
CORE
B3GNT5
PLEK
HLA-DQA1
CYBB
FPR1
OLR1
B3GNT5
IFI30
GPR183
EVI2B
PERIPHERAL
B3GNT5
PLEK
HLA-DQA1
CYBB
FPR1
OLR1
B3GNT5
IFI30
GPR183
EVI2B

### C8-Proliferative Myeloid



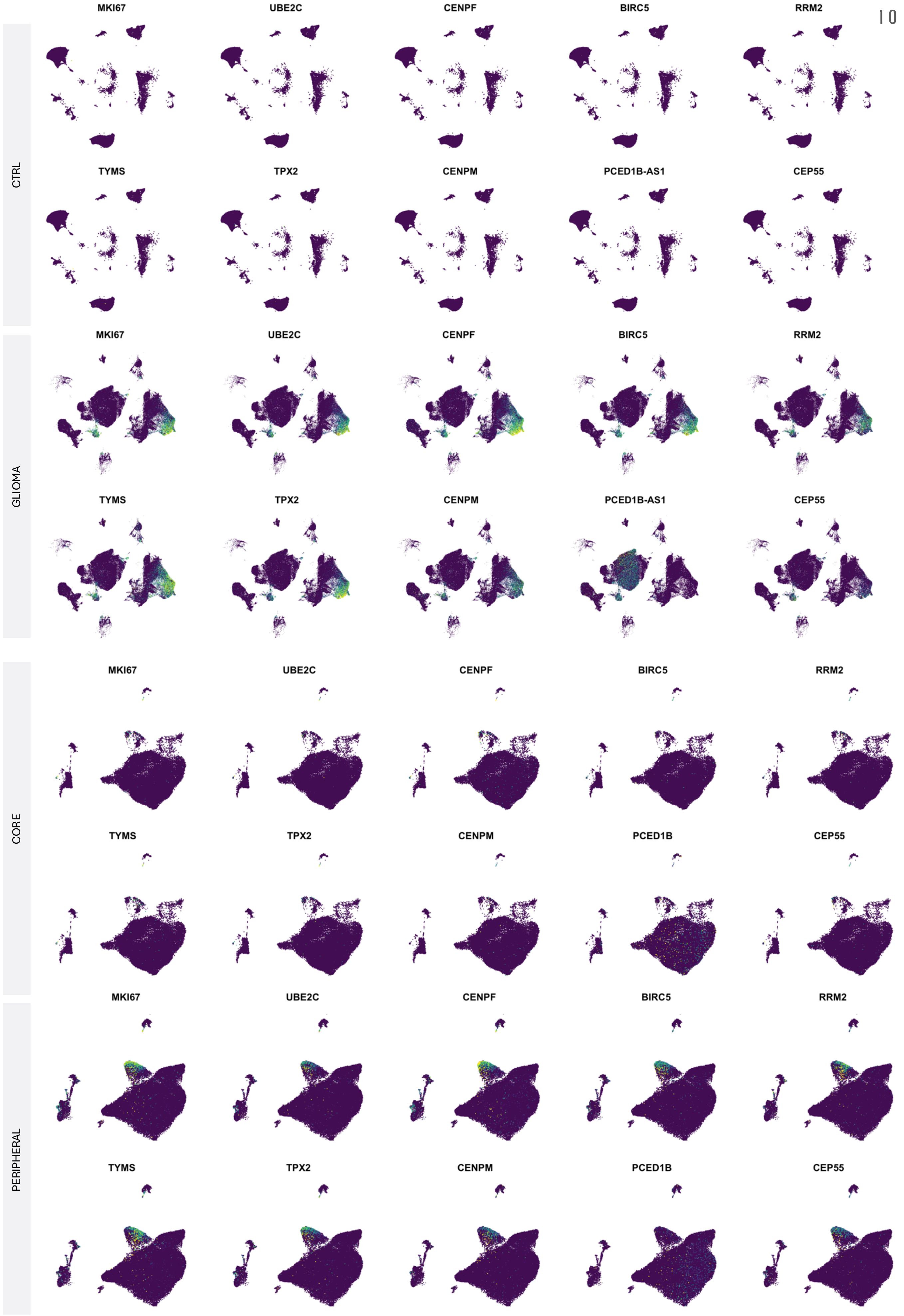

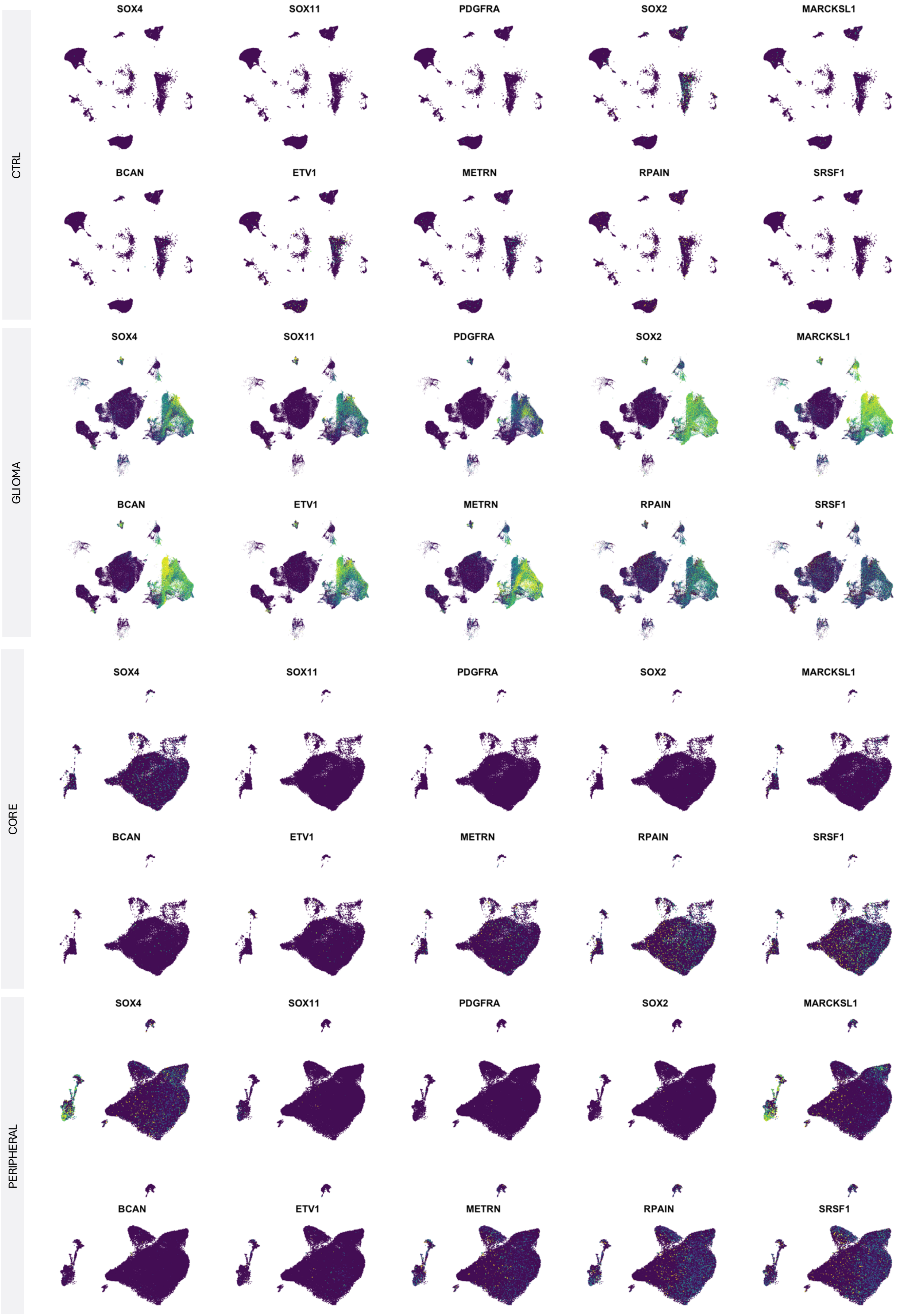
CTRL
SOX4
SOX11
PDGFRA
SOX2
MARCKSL1
BCAN
ETV1
METRN
RPAIN
SRSF1
GLIOMA
SOX4
SOX11
PDGFRA
SOX2
MARCKSL1
BCAN
ETV1
METRN
RPAIN
SRSF1
CORE
SOX4
SOX11
PDGFRA
SOX2
MARCKSL1
BCAN
ETV1
METRN
RPAIN
SRSF1
PERIPHERAL
SOX4
SOX11
PDGFRA
SOX2
MARCKSL1
BCAN
ETV1
METRN
RPAIN
SRSF1

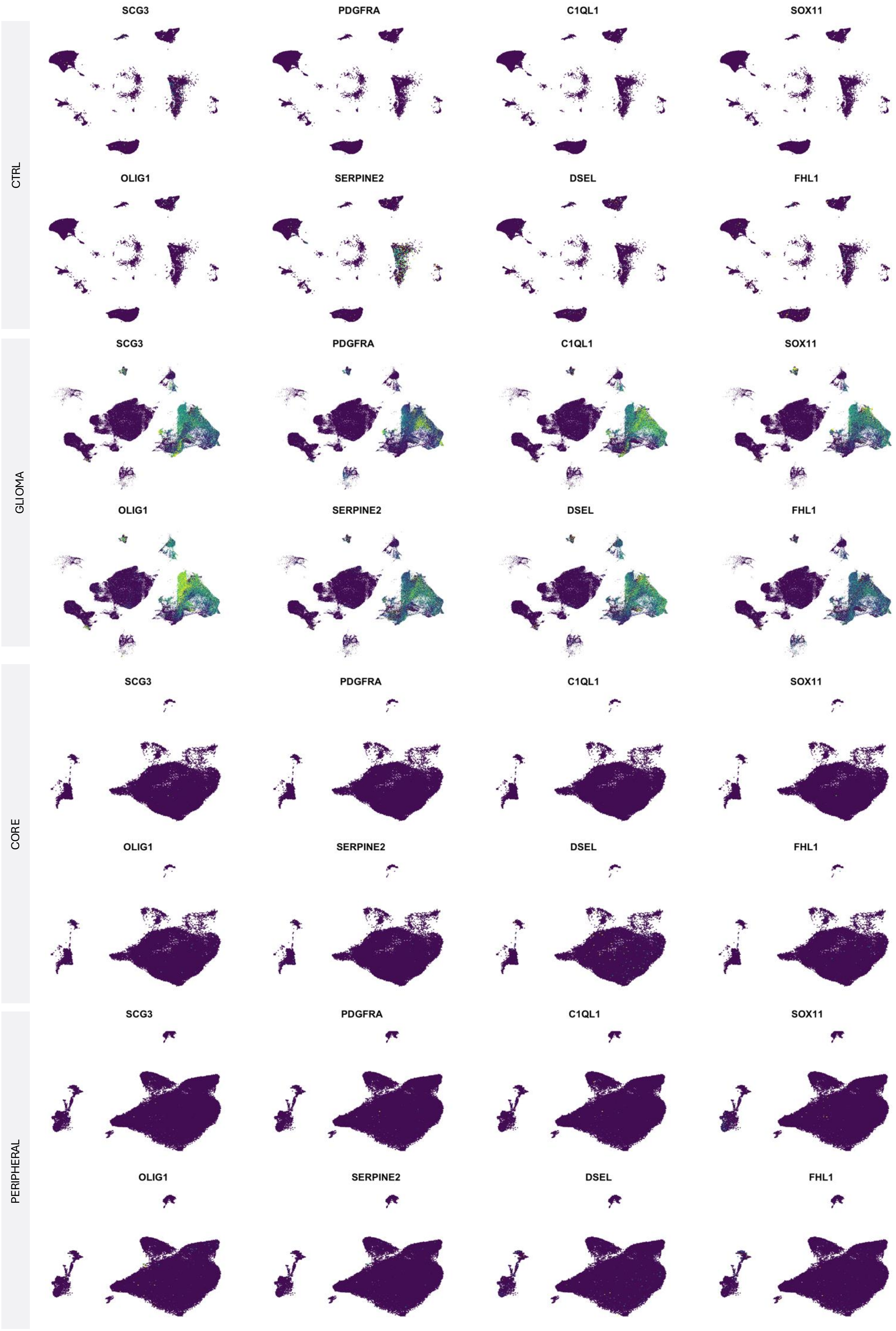
CTRL
SCG3
PDGFRA
C1QL1
SOX11
OLIG1
SERPINE2
DSEL
FHL1
GLIOMA
SCG3
PDGFRA
C1QL1
SOX11
OLIG1
SERPINE2
DSEL
FHL1
CORE
SCG3
PDGFRA
C1QL1
SOX11
OLIG1
SERPINE2
DSEL
FHL1
PERIPHERAL
SCG3
PDGFRA
C1QL1
SOX11
OLIG1
SERPINE2
DSEL
FHL1

### Analysis 2

Conservative Markers & Find All Markers

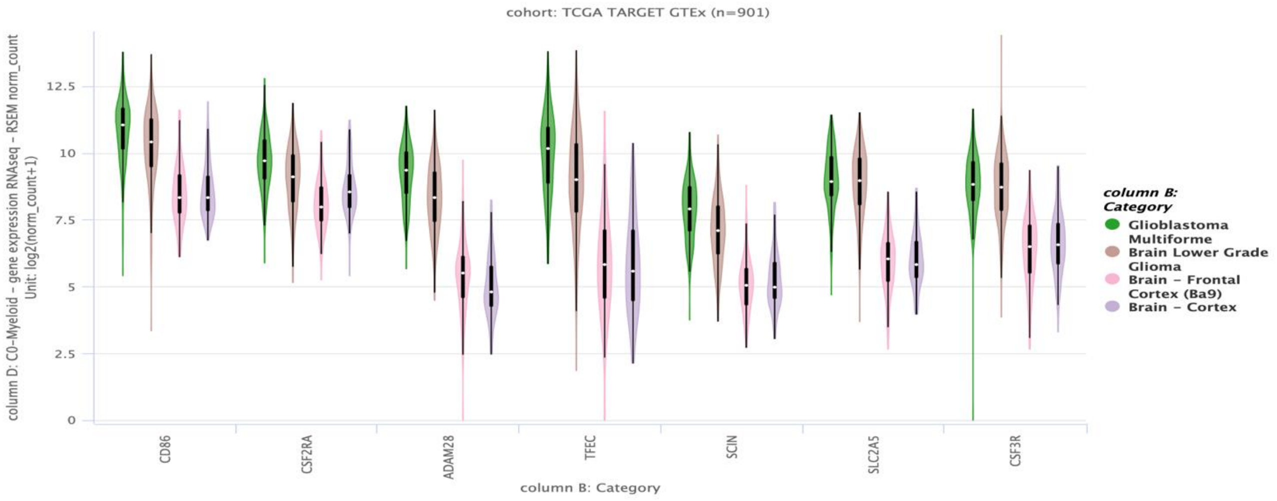
cohort: TCGA TARGET GTEx (n=901)
column D: C0-Myeloid – gene expression RNAseq – RSEM norm_count
Unit: log2(norm_count+1)
12.5
10
7.5
5
2.5
0
CD86
CSF2RA
ADAM28
TFEC
SCIN
SLC2A5
CSF3R
column B: Category
column B: Category
Glioblastoma Multiforme
Brain Lower Grade Glioma
Brain – Frontal Cortex (Ba9)
Brain – Cortex

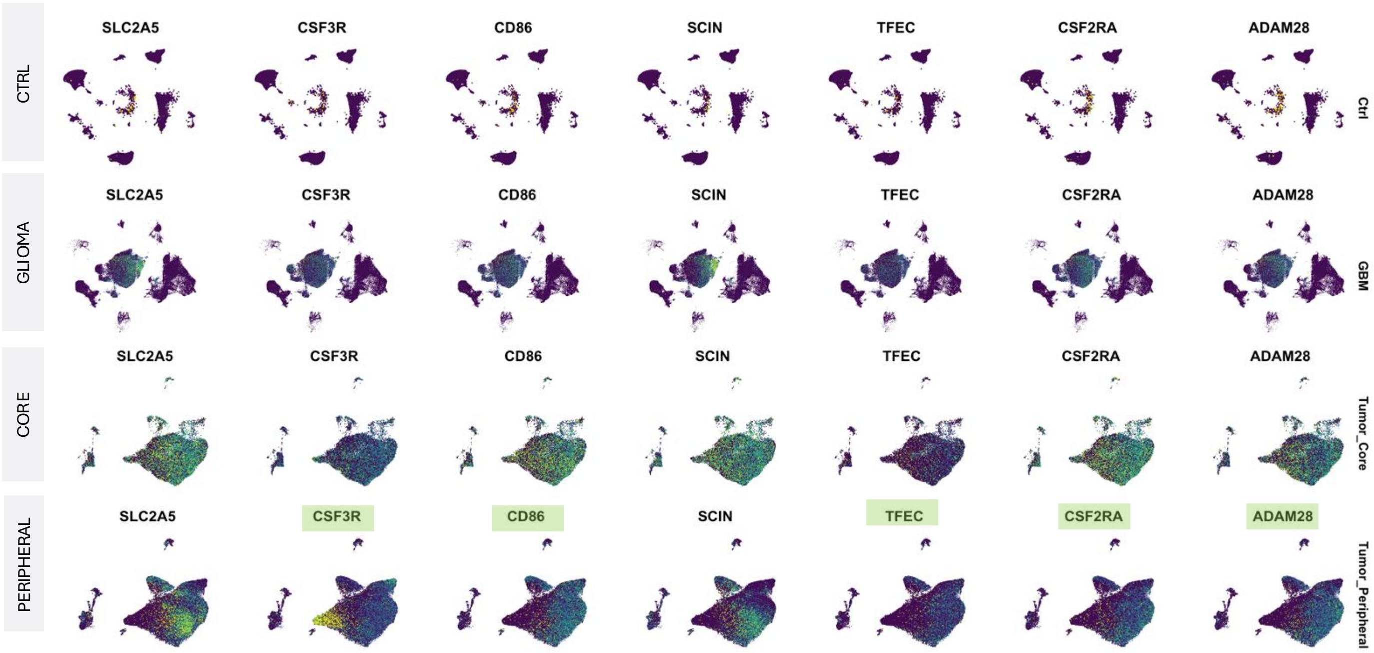
CTRL
GLIOMA
CORE
PERIPHERAL
SLC2A5
CSF3R
CD86
SCIN
TFEC
CSF2RA
ADAM28
Ctrl
GBM
Tumor_Core
Tumor_Peripheral

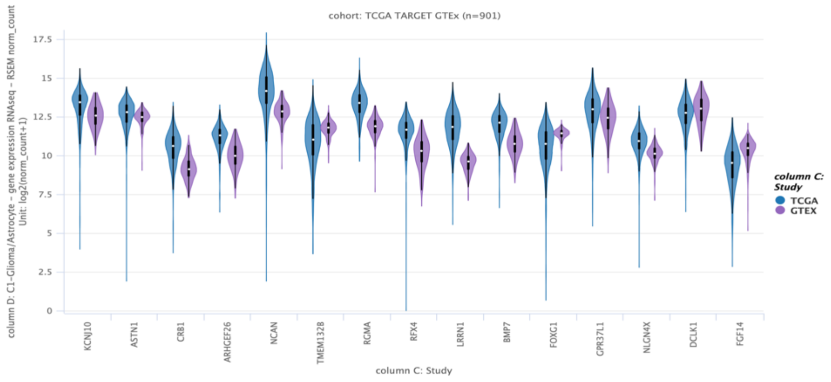
cohort: TCGA TARGET GTEx (n=901)
column D: C1-Glioma/Astrocyte - gene expression RNAseq - RSEM norm_count
Unit: log2(norm_count+1)
17.5
15
12.5
10
7.5
5
2.5
0
KCNJ10
ASTN1
CRB1
ARHGEF26
NCAN
TMEM132B
RGMA
RFX4
LRRN1
BMP7
FOXG1
GPR37L1
NLGN4X
DCLK1
FGF14
column C: Study
column C: Study
TCGA
GTEX

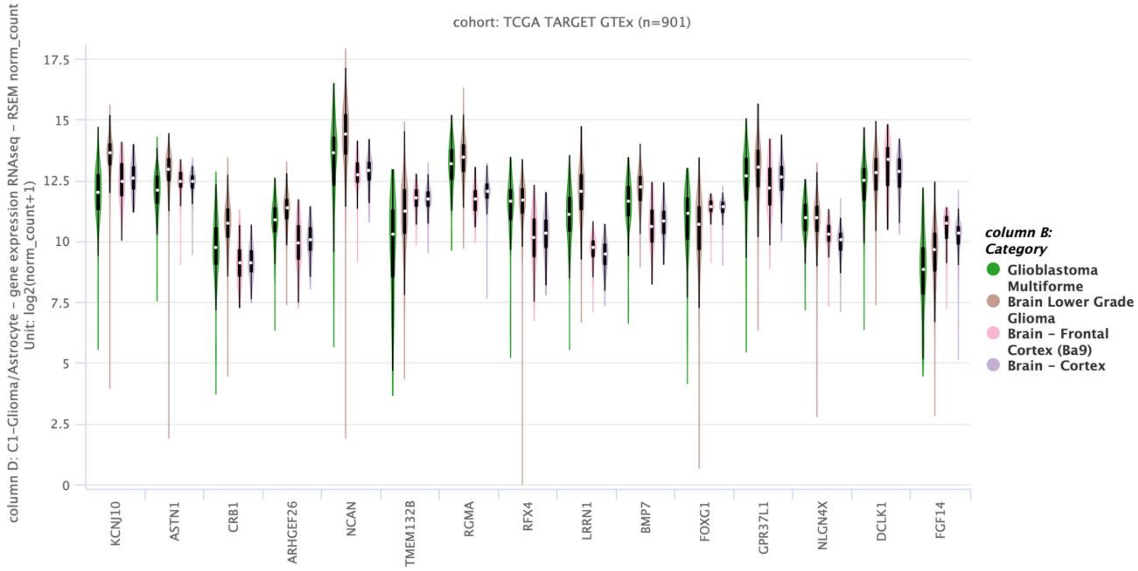
cohort: TCGA TARGET GTEx (n=901)
column D: C1-Glioma/Astrocyte - gene expression RNAseq - RSEM norm_count
Unit: log2(norm_count+1)
17.5
15
12.5
10
7.5
5
2.5
0
KCNJ10
ASTN1
CRB1
ARHGEF26
NCAN
TMEM132B
RGMA
RFX4
LRRN1
BMP7
FOXG1
GPR37L1
NLGN4X
DCLK1
FGF14
column B: Category
Glioblastoma Multiforme
Brain Lower Grade Glioma
Brain - Frontal Cortex (Ba9)
Brain - Cortex

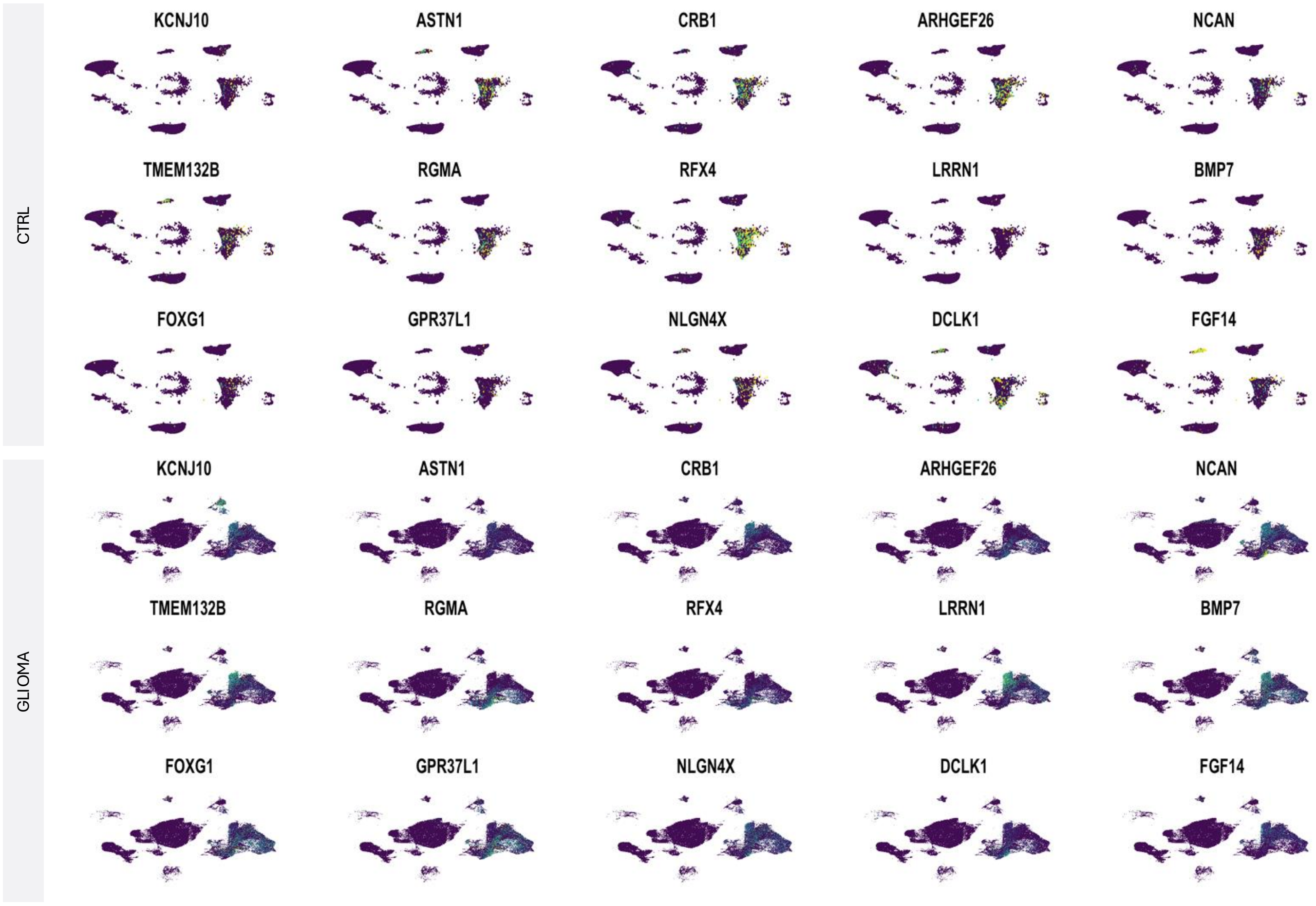
CTRL
GLIOMA
KCNJ10
ASTN1
CRB1
ARHGEF26
NCAN
TMEM132B
RGMA
RFX4
LRRN1
BMP7
FOXG1
GPR37L1
NLGN4X
DCLK1
FGF14

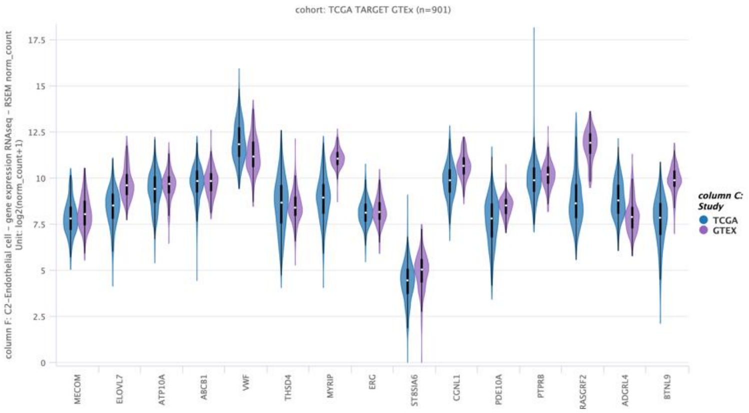
cohort: TCGA TARGET GTEx (n=901)
column F: C2-Endothelial cell - gene expression RNAseq - RSEM norm_count
Unit: log2(norm_count+1)
17.5
15
12.5
10
7.5
5
2.5
0
MECOM
ELOVL7
ATP10A
ABCB1
VWF
THSD4
MYRIP
ERG
ST8SIA6
CGNL1
PDE10A
PTPRB
RASGRF2
ADGRL4
BTNL9
column C: Study
TCGA
GTEX

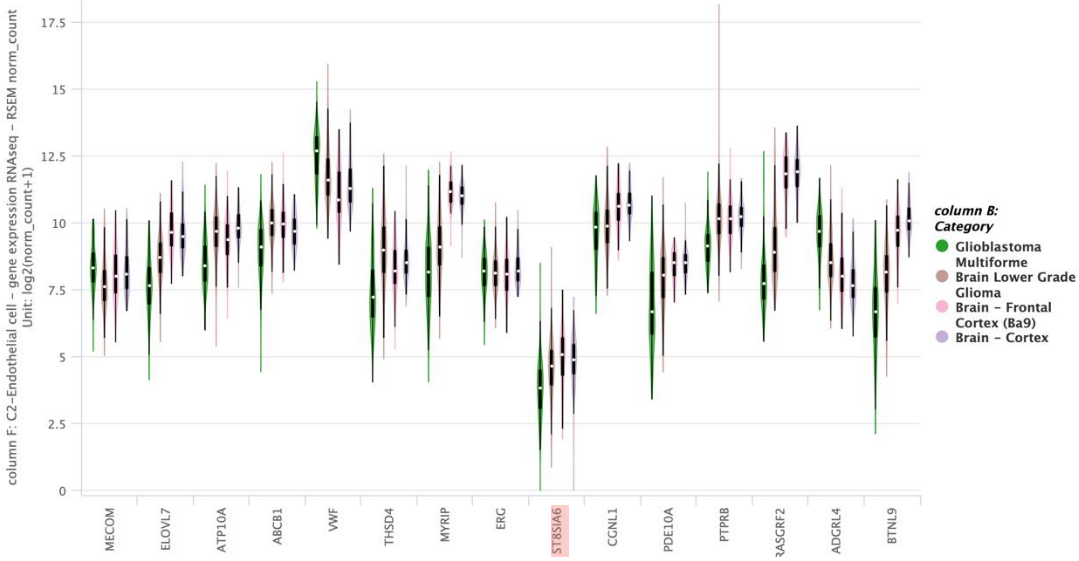
column F: C2-Endothelial cell – gene expression RNAseq – RSEM norm_count
Unit: log2(norm_count+1)
17.5
15
12.5
10
7.5
5
2.5
0
MECOM
ELOVL7
ATP10A
ABCB1
VWF
THSD4
MYRIP
ERG
ST8SIA6
CGNL1
PDE10A
PTPRB
RASGRF2
ADGRL4
BTNL9
column B: Category
Glioblastoma Multiforme
Brain Lower Grade Glioma
Brain – Frontal Cortex (Ba9)
Brain – Cortex

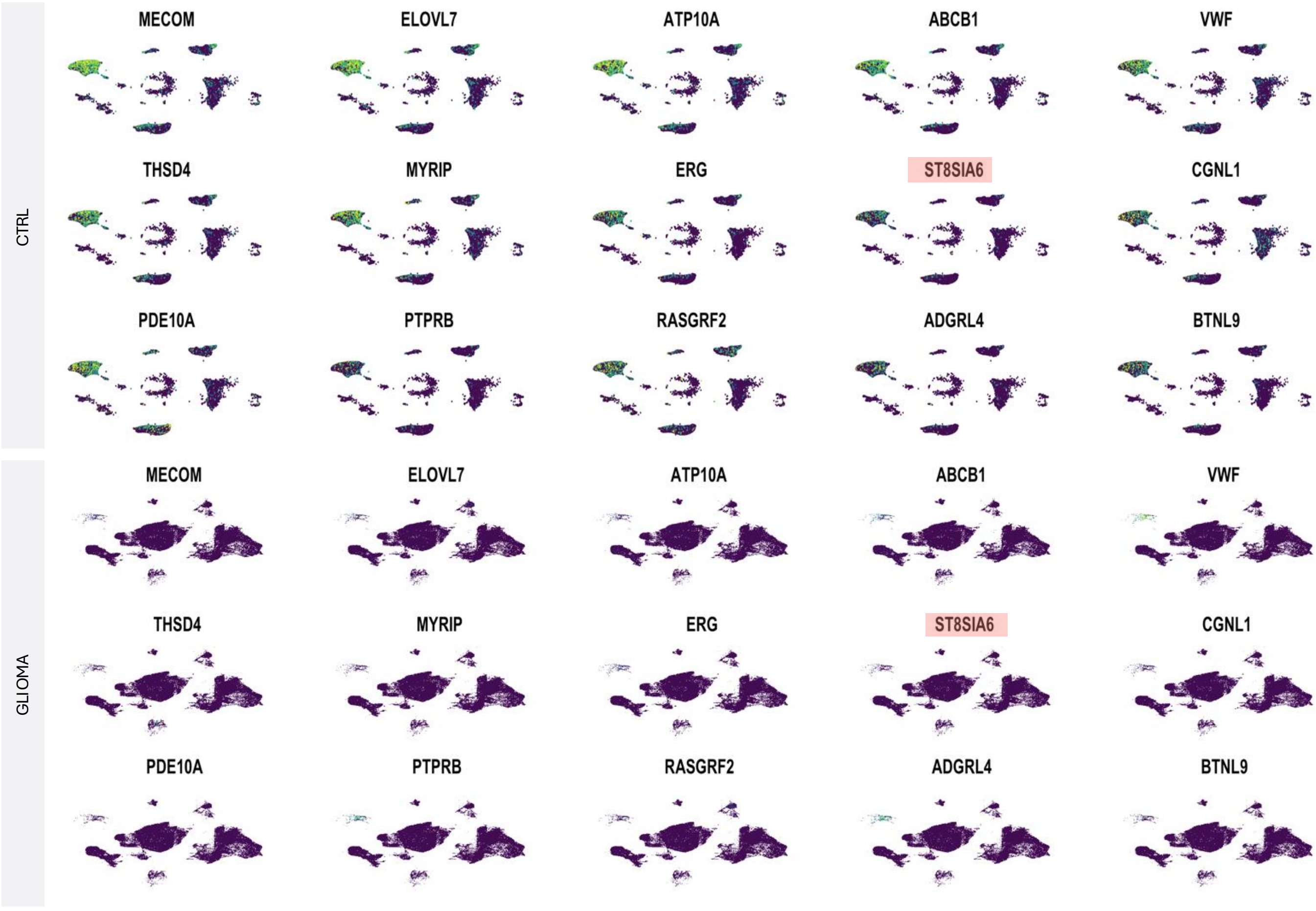
CTRL
MECOM
ELOVL7
ATP10A
ABCB1
VWF
THSD4
MYRIP
ERG
ST8SIA6
CGNL1
PDE10A
PTPRB
RASGRF2
ADGRL4
BTNL9
GLIOMA
MECOM
ELOVL7
ATP10A
ABCB1
VWF
THSD4
MYRIP
ERG
ST8SIA6
CGNL1
PDE10A
PTPRB
RASGRF2
ADGRL4
BTNL9

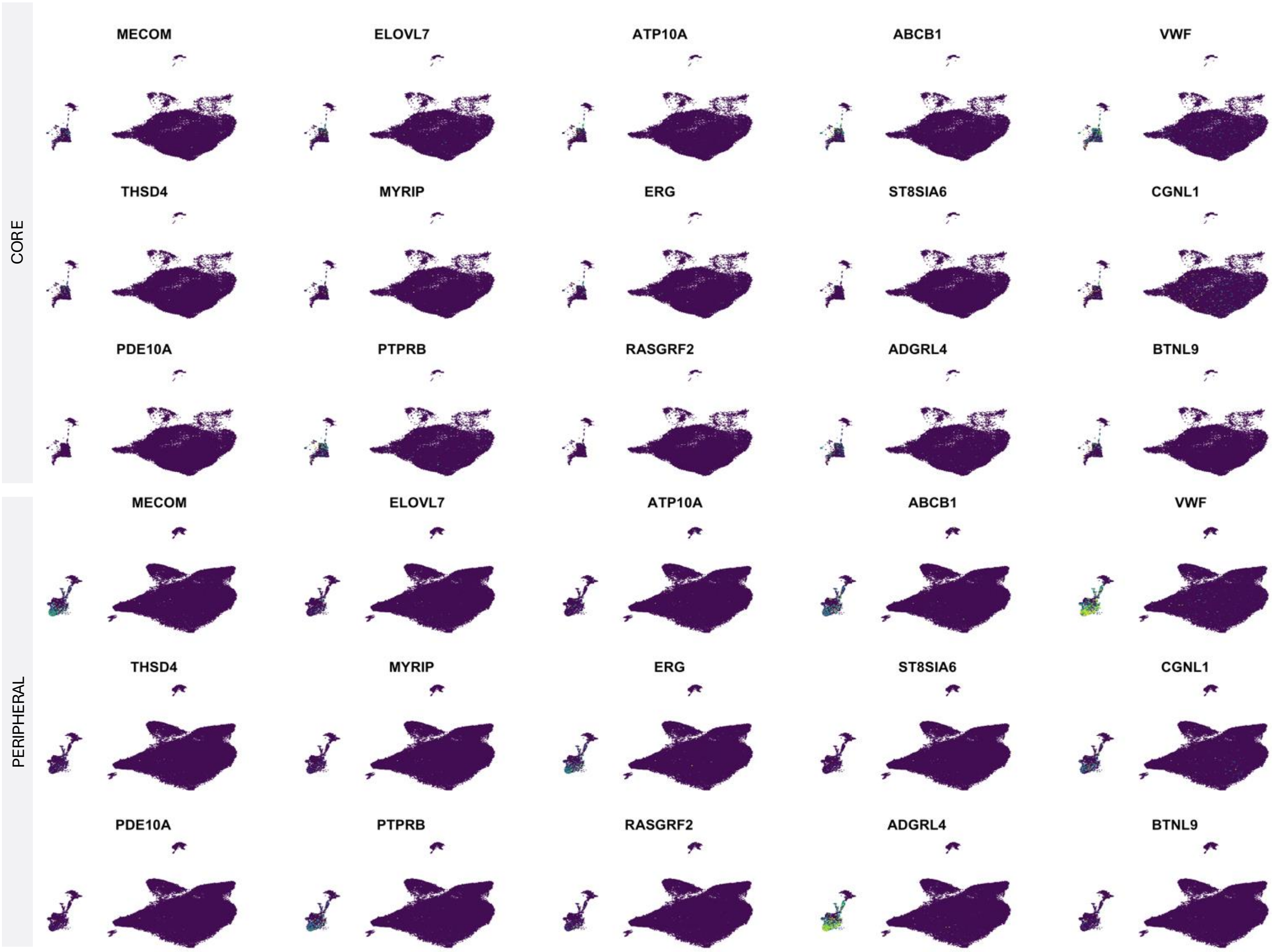
CORE
MECOM
ELOVL7
ATP10A
ABCB1
VWF
THSD4
MYRIP
ERG
ST8SIA6
CGNL1
PDE10A
PTPRB
RASGRF2
ADGRL4
BTNL9
PERIPHERAL
MECOM
ELOVL7
ATP10A
ABCB1
VWF
THSD4
MYRIP
ERG
ST8SIA6
CGNL1
PDE10A
PTPRB
RASGRF2
ADGRL4
BTNL9

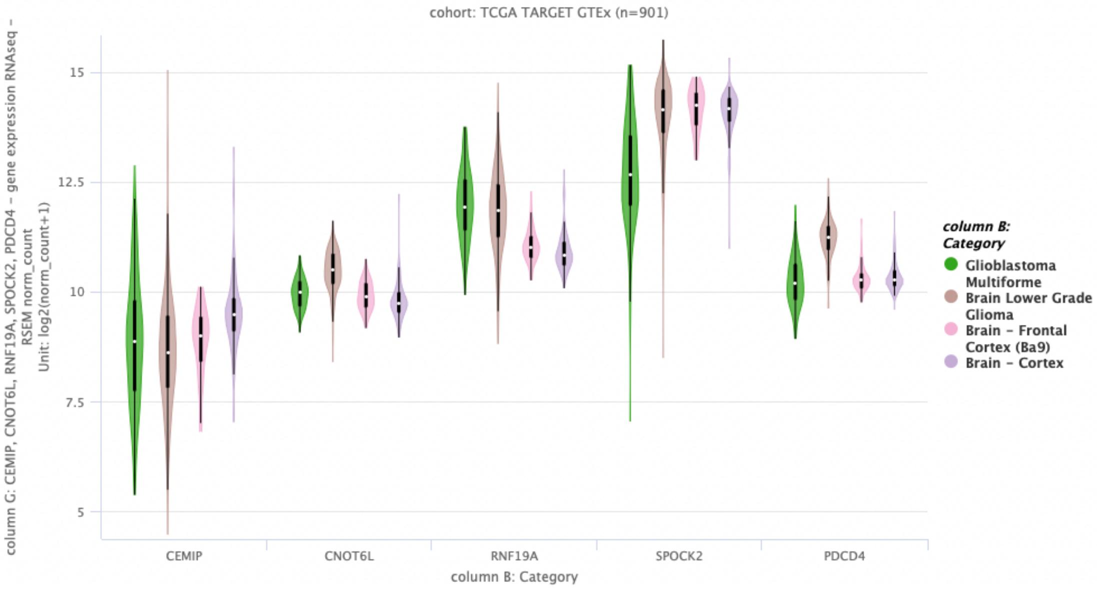
cohort: TCGA TARGET GTEx (n=901)
column G: CEMIP, CNOT6L, RNF19A, SPOCK2, PDCD4 - gene expression RNAseq - RSEM norm_count
Unit: log2(norm_count+1)
15
12.5
10
7.5
5
CEMIP
CNOT6L
RNF19A
SPOCK2
PDCD4
column B: Category
column B: Category
Glioblastoma Multiforme
Brain Lower Grade Glioma
Brain - Frontal Cortex (Ba9)
Brain - Cortex

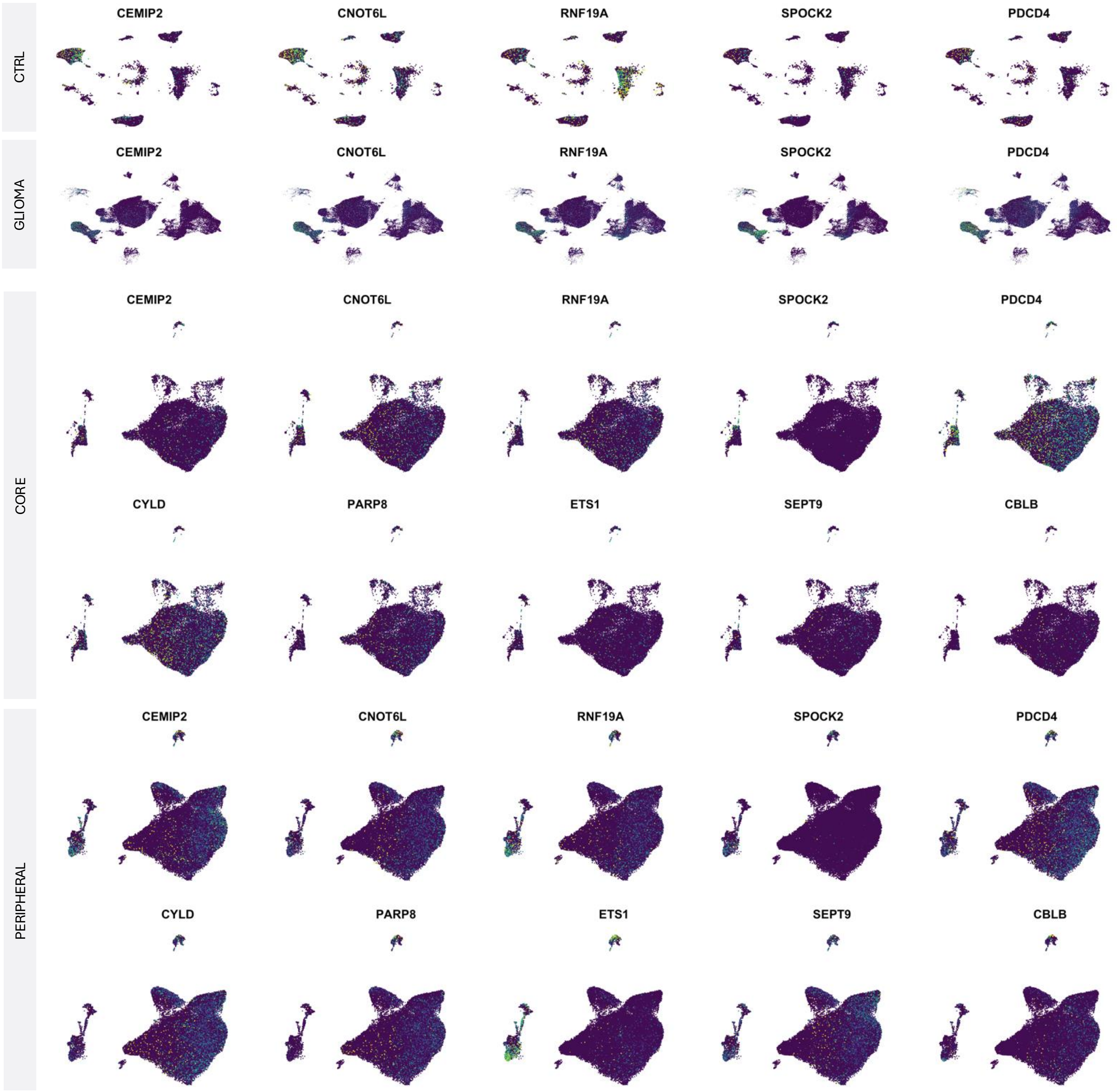
CTRL
GLIOMA
CORE
PERIPHERAL
CEMIP2
CNOT6L
RNF19A
SPOCK2
PDCD4
CYLD
PARP8
ETS1
SEPT9
CBLB

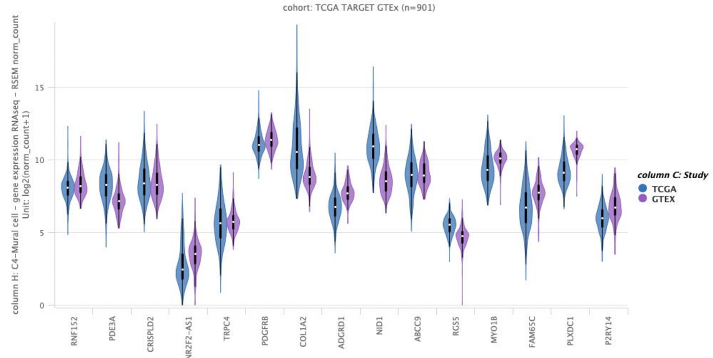
cohort: TCGA TARGET GTEx (n=901)
column H: C4-Mural cell – gene expression RNAseq – RSEM norm_count
Unit: log2(norm_count+1)
column C: Study
TCGA
GTEX
RNF152
PDE3A
CRISPLD2
NR2F2-AS1
TRPC4
PDGFRB
COL1A2
ADGRD1
NID1
ABCC9
RGS5
MYO1B
FAM65C
PLXDC1
P2RY14

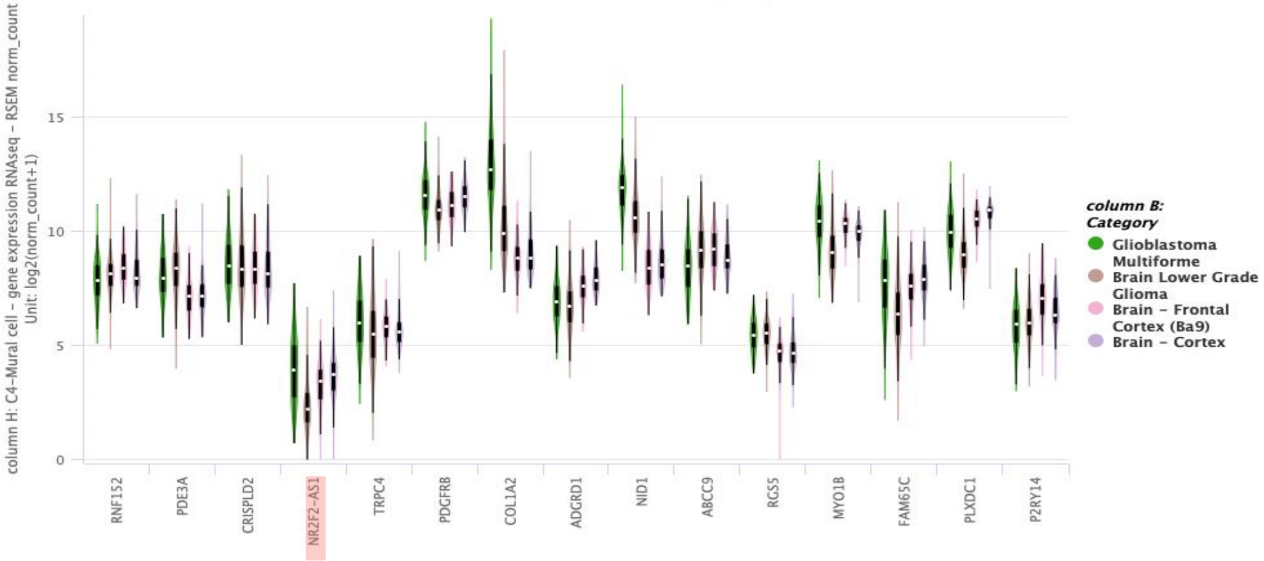
column H: C4-Mural cell – gene expression RNAseq – RSEM norm_count
Unit: log2(norm_count+1)
column B: Category
Glioblastoma Multiforme
Brain Lower Grade Glioma
Brain – Frontal Cortex (Ba9)
Brain – Cortex
RNF152
PDE3A
CRISPLD2
NR2F2-AS1
TRPC4
PDGFRB
COL1A2
ADGRD1
NID1
ABCC9
RGS5
MYO1B
FAM65C
PLXDC1
P2RY14

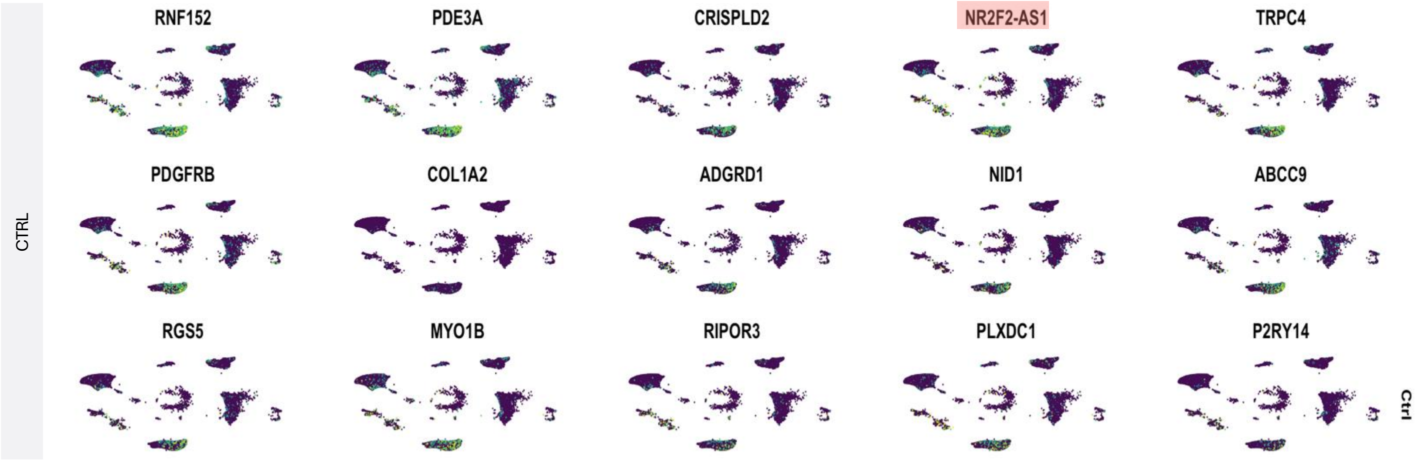
CTRL
RNF152
PDE3A
CRISPLD2
NR2F2-AS1
TRPC4
PDGFRB
COL1A2
ADGRD1
NID1
ABCC9
RGS5
MYO1B
RIPOR3
PLXDC1
P2RY14
Ctrl

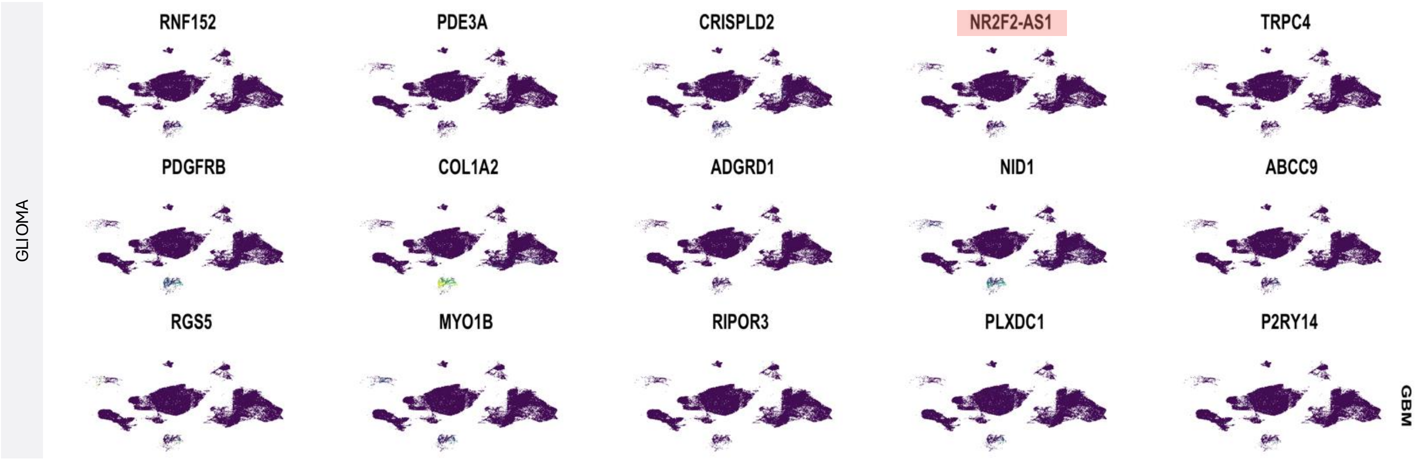
GLIOMA
RNF152
PDE3A
CRISPLD2
NR2F2-AS1
TRPC4
PDGFRB
COL1A2
ADGRD1
NID1
ABCC9
RGS5
MYO1B
RIPOR3
PLXDC1
P2RY14
GBM

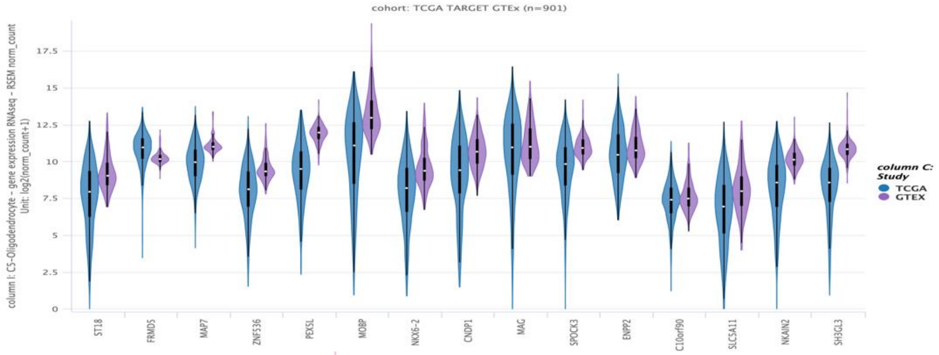
cohort: TCGA TARGET GTEx (n=901)
column I: C5-Oligodendrocyte - gene expression RNAseq - RSEM norm_count
Unit: log2(norm_count+1)
column C: Study
TCGA
GTEX
ST18
FRMD5
MAP7
ZNF536
PEX5L
MOBP
NKX6-2
CNDP1
MAG
SPOCK3
ENPP2
C10orf90
SLC5A11
NKAIN2
SH3GL3

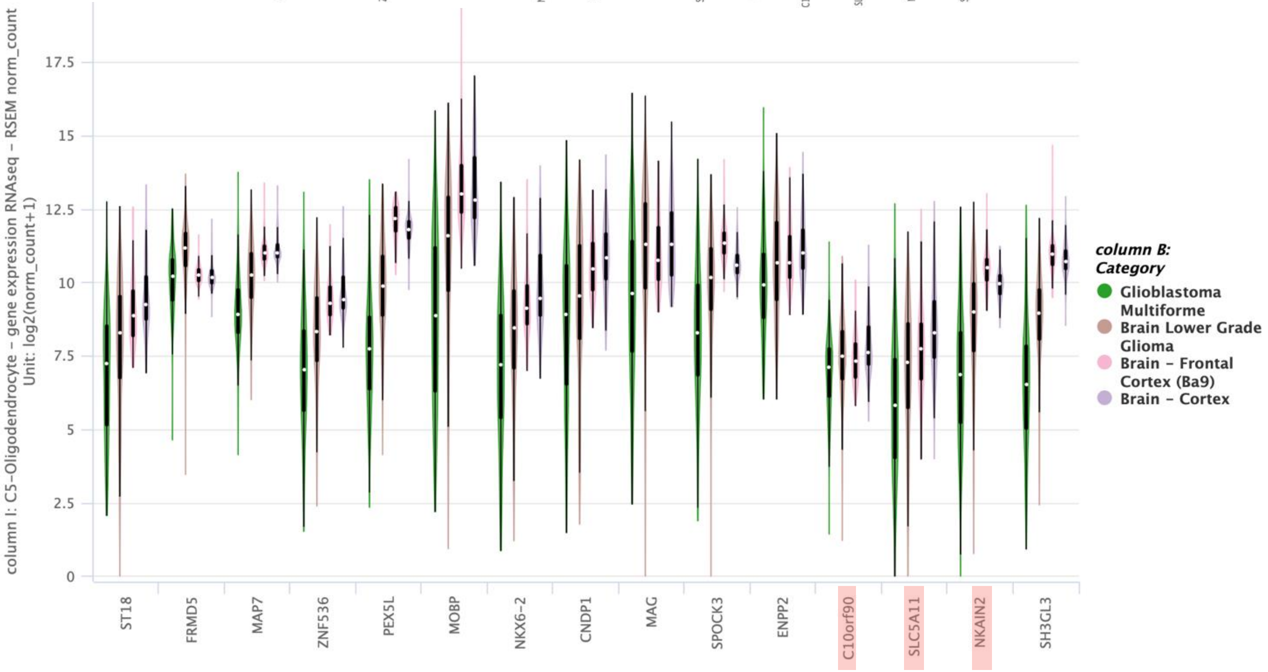
column I: C5-Oligodendrocyte - gene expression RNAseq - RSEM norm_count
Unit: log2(norm_count+1)
column B: Category
Glioblastoma Multiforme
Brain Lower Grade Glioma
Brain - Frontal Cortex (Ba9)
Brain - Cortex
ST18
FRMD5
MAP7
ZNF536
PEX5L
MOBP
NKX6-2
CNDP1
MAG
SPOCK3
ENPP2
C10orf90
SLC5A11
NKAIN2
SH3GL3

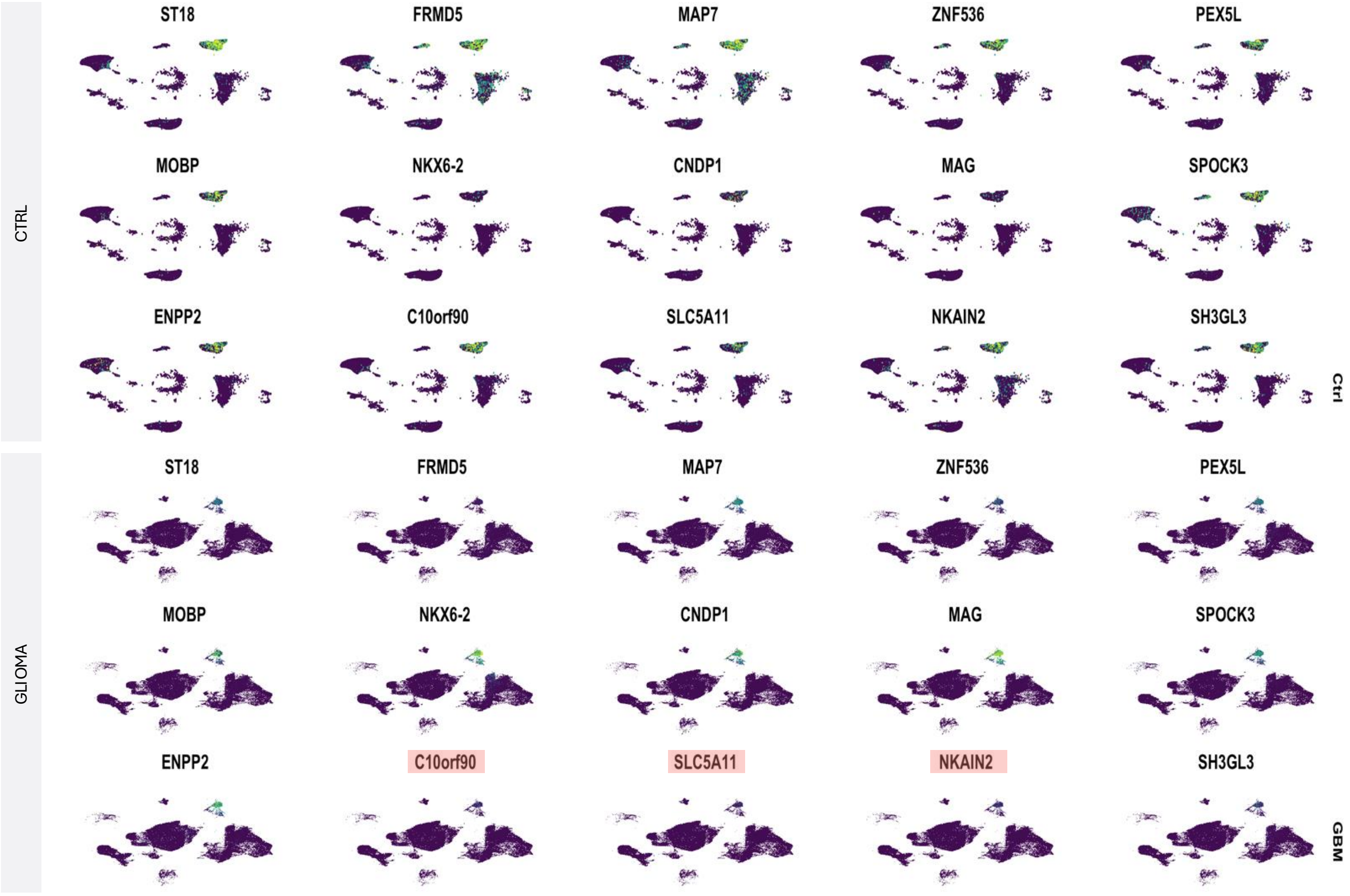
CTRL
ST18
FRMD5
MAP7
ZNF536
PEX5L
MOBP
NKX6-2
CNDP1
MAG
SPOCK3
ENPP2
C10orf90
SLC5A11
NKAIN2
SH3GL3
Ctrl
GLIOMA
ST18
FRMD5
MAP7
ZNF536
PEX5L
MOBP
NKX6-2
CNDP1
MAG
SPOCK3
ENPP2
C10orf90
SLC5A11
NKAIN2
SH3GL3
GBM

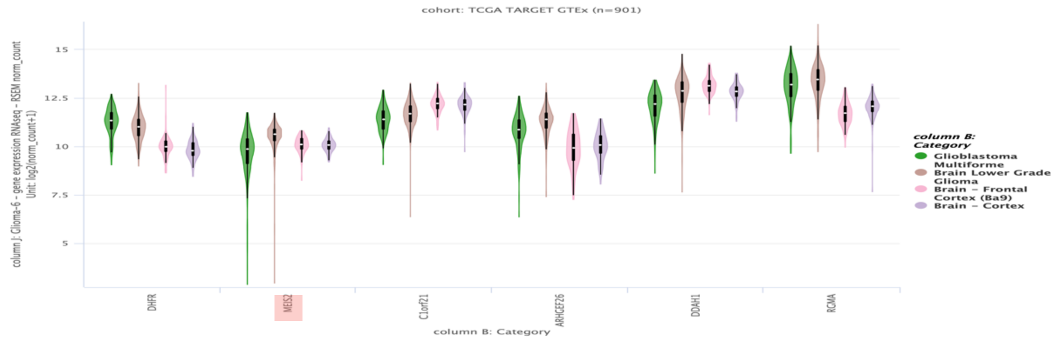
cohort: TCGA TARGET GTEx (n=901)
column J: Glioma-6 - gene expression RNAseq - RSEM norm_count
Unit: log2(norm_count+1)
15
12.5
10
7.5
5
DHFR
MEIS2
C1orf21
ARHGEF26
DDAH1
RGMA
column B: Category
column B: Category
Glioblastoma Multiforme
Brain Lower Grade Glioma
Brain - Frontal Cortex (Ba9)
Brain - Cortex

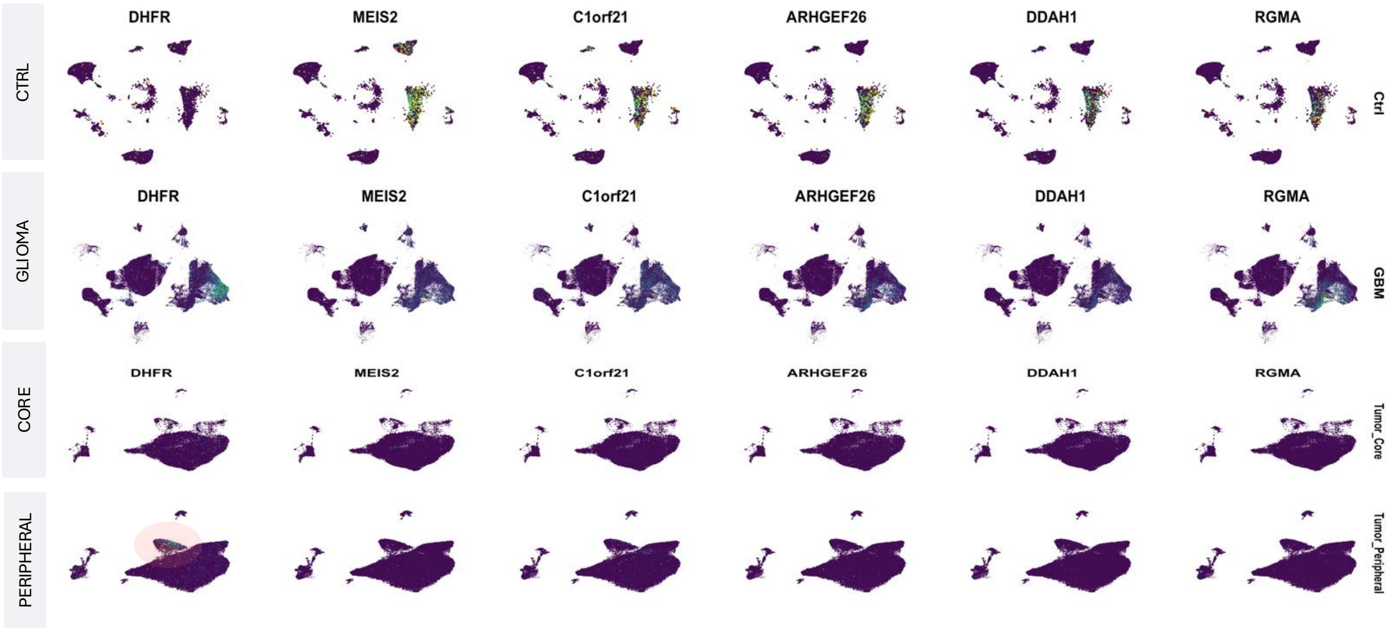
CTRL
GLIOMA
CORE
PERIPHERAL
DHFR
MEIS2
C1orf21
ARHGEF26
DDAH1
RGMA
Ctrl
GBM
Tumor_Core
Tumor_Peripheral

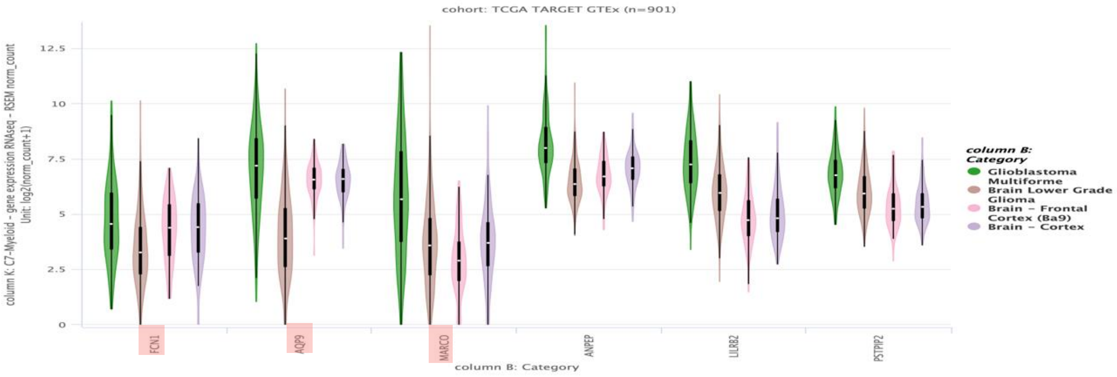
cohort: TCGA TARGET GTEx (n=901)
column K: C7-Myeloid - gene expression RNAseq - RSEM norm_count
Unit: log2(norm_count+1)
12.5
10
7.5
5
2.5
0
FCN1
AQP9
MARCO
ANPEP
LILRB2
PSTPIP2
column B: Category
column B: Category
Glioblastoma Multiforme
Brain Lower Grade Glioma
Brain - Frontal Cortex (Ba9)
Brain - Cortex

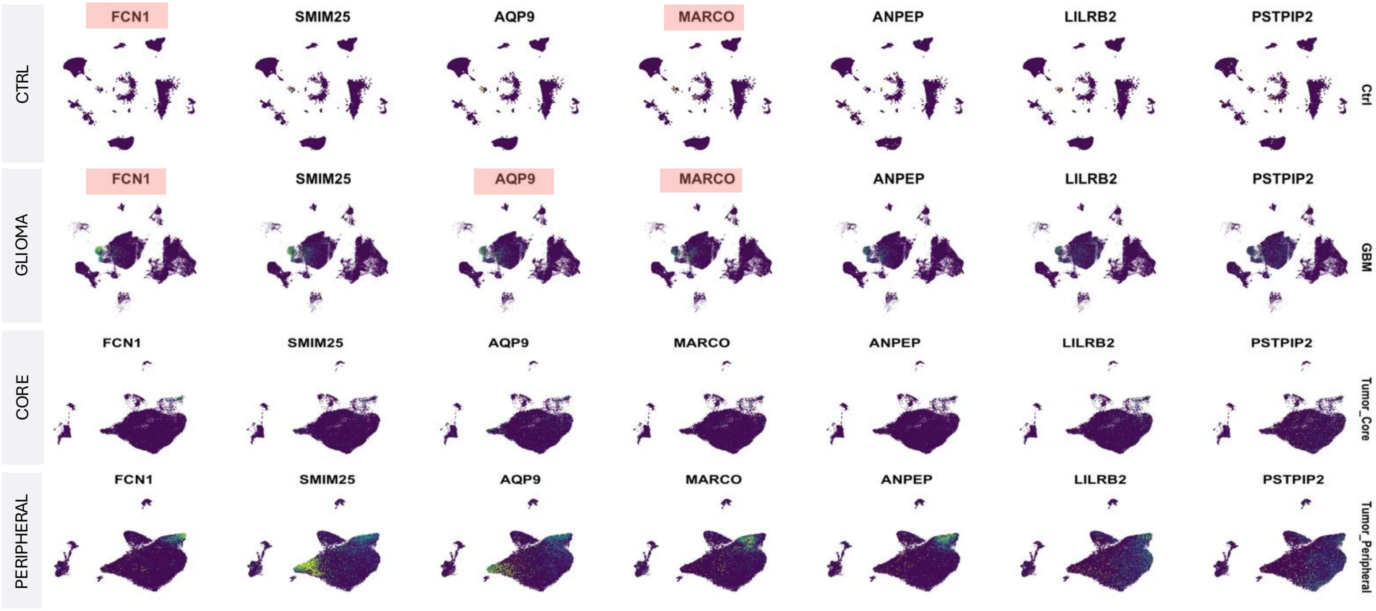
CTRL
GLIOMA
CORE
PERIPHERAL
FCN1
SMIM25
AQP9
MARCO
ANPEP
LILRB2
PSTPIP2
Ctrl
GBM
Tumor_Core
Tumor_Peripheral

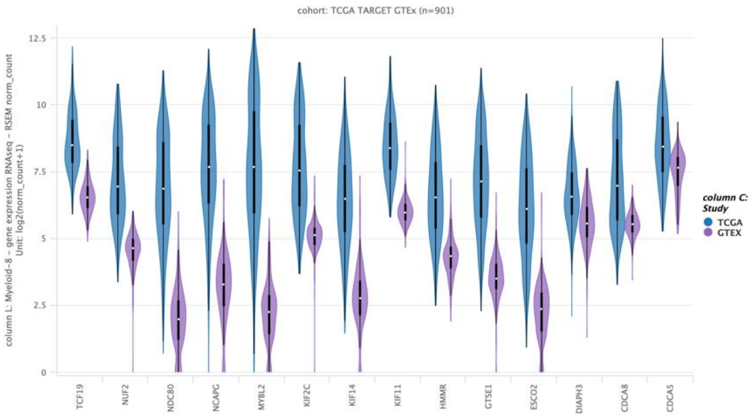
cohort: TCGA TARGET GTEx (n=901)
column L: Myeloid-8 - gene expression RNAseq - RSEM norm_count
Unit: log2(norm_count+1)
12.5
10
7.5
5
2.5
0
TCF19
NUF2
NDC80
NCAPG
MYBL2
KIF2C
KIF14
KIF11
HMMR
GTSE1
ESCO2
DIAPH3
CDCA8
CDCA5
column C: Study
TCGA
GTEX

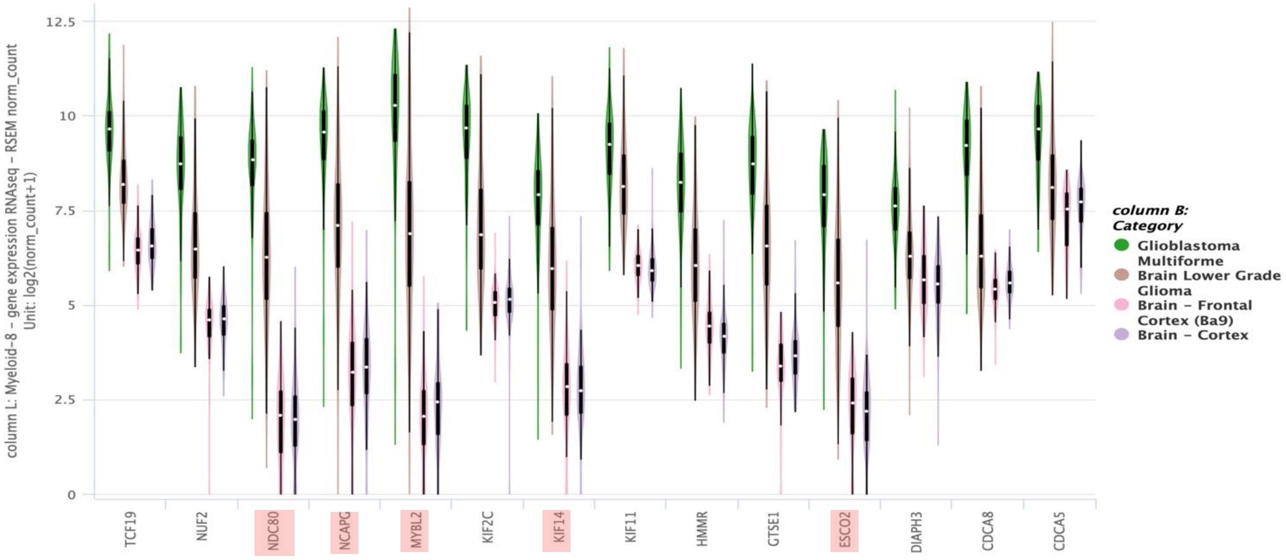
column L: Myeloid-8 - gene expression RNAseq - RSEM norm_count
Unit: log2(norm_count+1)
12.5
10
7.5
5
2.5
0
TCF19
NUF2
NDC80
NCAPG
MYBL2
KIF2C
KIF14
KIF11
HMMR
GTSE1
ESCO2
DIAPH3
CDCA8
CDCA5
column B: Category
Glioblastoma Multiforme
Brain Lower Grade Glioma
Brain - Frontal Cortex (Ba9)
Brain - Cortex

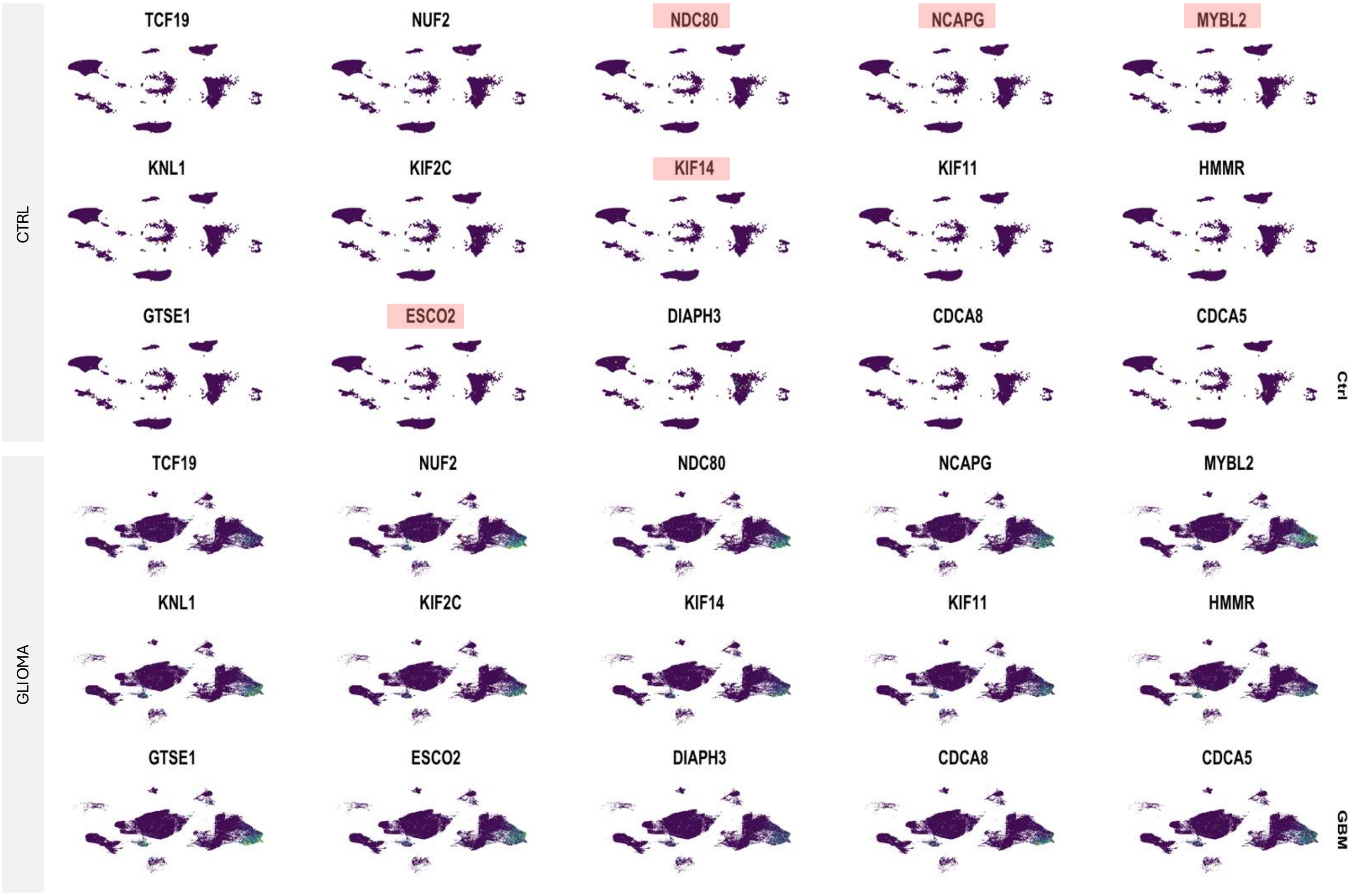
CTRL
TCF19
NUF2
NDC80
NCAPG
MYBL2
KNL1
KIF2C
KIF14
KIF11
HMMR
GTSE1
ESCO2
DIAPH3
CDCA8
CDCA5
Ctrl
GLIOMA
TCF19
NUF2
NDC80
NCAPG
MYBL2
KNL1
KIF2C
KIF14
KIF11
HMMR
GTSE1
ESCO2
DIAPH3
CDCA8
CDCA5
GBM

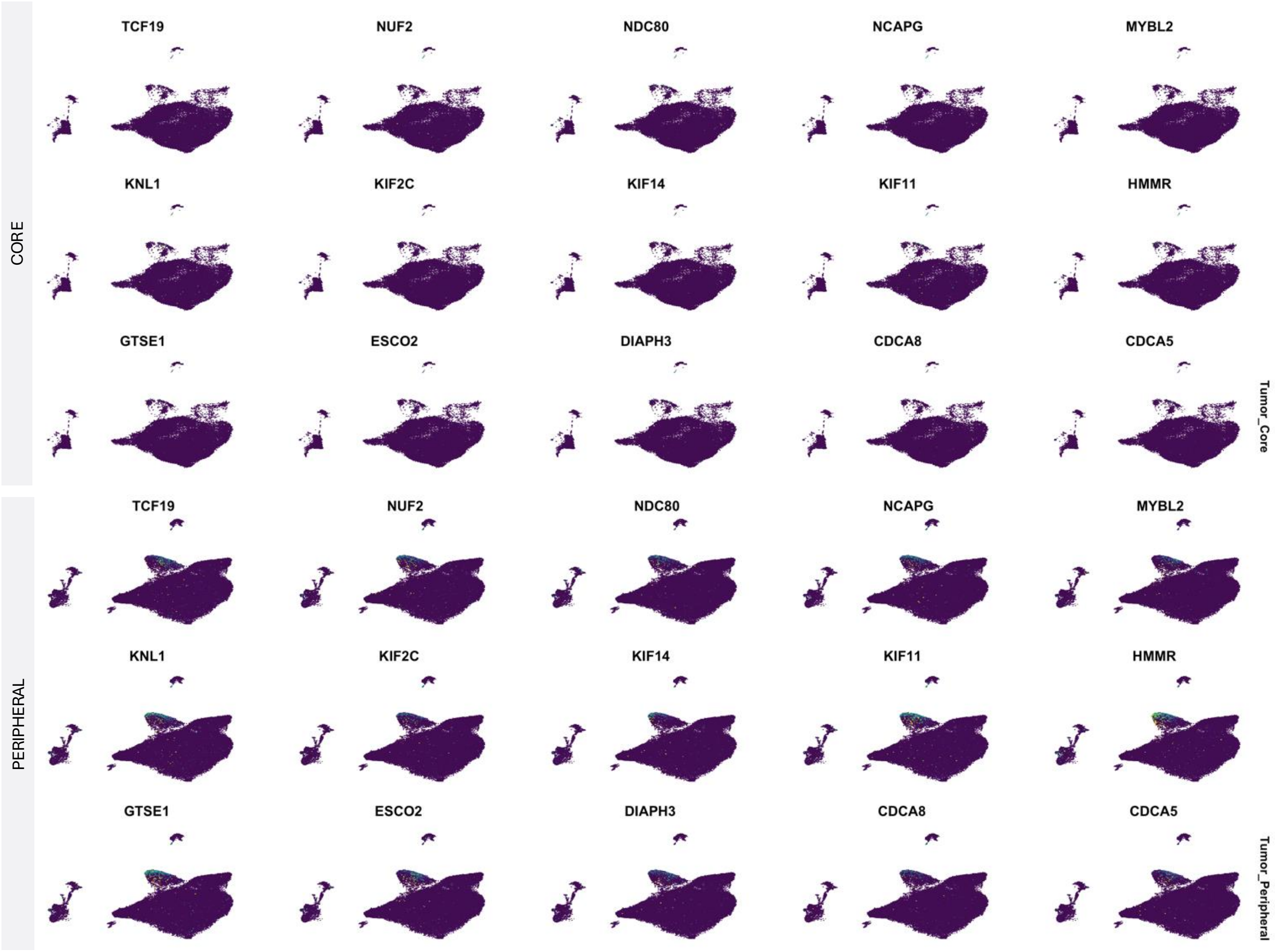
CORE
TCF19
NUF2
NDC80
NCAPG
MYBL2
KNL1
KIF2C
KIF14
KIF11
HMMR
GTSE1
ESCO2
DIAPH3
CDCA8
CDCA5
Tumor_Core
PERIPHERAL
TCF19
NUF2
NDC80
NCAPG
MYBL2
KNL1
KIF2C
KIF14
KIF11
HMMR
GTSE1
ESCO2
DIAPH3
CDCA8
CDCA5
Tumor_Peripheral
